# CENP-C unwraps the CENP-A nucleosome through the H2A C-terminal tail

**DOI:** 10.1101/705699

**Authors:** Ahmad Ali-Ahmad, Silvija Bilokapić, Ingmar B. Schäfer, Mario Halić, Nikolina Sekulić

## Abstract

Centromeres are defined epigenetically by nucleosomes containing the histone H3 variant CENP-A, upon which the constitutive centromere-associated network of proteins (CCAN) is built. CENP-C, is considered to be a central organizer of the CCAN. We provide new molecular insights into the structure of CENP-A nucleosomes, in isolation and in complex with the CENP-C central region (CENP-C^CR^), the main CENP-A binding module of CENP-C. We establish that the short αN-helix of CENP-A promotes DNA flexibility at the nucleosome ends, independently of the sequence it wraps.

Furthermore, we show that, in vitro, two regions of CENP-C (CENP-C^CR^ and CENP-C^motif^) both bind exclusively to the CENP-A nucleosome. We find CENP-C^CR^ to bind with high affinity due to an extended hydrophobic area made up of CENP-A^V532^ and CENP-A^V533^. Importantly, we identify two key conformational changes within the CENP-A nucleosome upon CENP-C binding. First, the loose DNA wrapping of CENP-A nucleosomes is further exacerbated, through destabilization of the H2A N-terminal tail. Second, CENP-C^CR^ rigidifies the N-terminal tail of H4 in the conformation favoring H4^K20^ monomethylation, essential for a functional centromere.

**Synopsis:** CENP-A nucleosomes have a short αN helix incompatible with complete DNA wrapping, independently of DNA sequence. CENP-C binds exclusively to CENP-A nucleosomes and this binding induces conformational changes that further differentiate CENP-A-containing from canonical nucleosomes.

- CENP-C binds CENP-A nucleosomes specifically
- DNA ends of the CENP-A nucleosome are further unwrapped in the CENP-A/CENP-C complex, due to flexible H2A C-terminal tails
- The N-terminal tail of H4 adopts a conformation favored for centromere specific H4^K20^ monomethylation when CENP-C is bound

## Introduction

The centromere is a chromosomal locus that directs accurate segregation of chromosomes during cell division [1]. Defects in chromosome segregation lead to aneuploidy, a hallmark of cancer [2].

DNA sequences underlining human centromeres are composed of AT rich repeats (termed α-satellites), but these are neither necessary nor sufficient for centromere function. Instead, centromeres are specified epigenetically, by the presence of the histone H3 variant, CENP-A [reviewed in [3]]. Chromatin containing CENP-A nucleosomes must have unique structural properties to organize the constitutive centromere associated network (CCAN). It is therefore critical to gain a full structural understanding of the CENP-A nucleosome alone and in complex with the two key components of the CCAN with which it interacts directly and specifically – CENP-C and CENP-N [4]. Initial clues on CENP-A nucleosome-specific features came from crystallographic studies of the (CENP-A/H4)_2_ tetramer [5] and of the CENP-A nucleosome [6]. These studies implied that the CENP-A nucleosome has an octameric histone core, similar to canonical nucleosomes in composition and structure. DNA ends in the crystal structure of CENP-A nucleosome [6] are disordered, indicating increased DNA flexibility. Recently, cryoEM structures of the human CENP-A nucleosome in complex with human CENP-N have been reported by several groups [7–9], revealing high-resolution molecular determinants for the CENP-A/CENP-N interaction.

CENP-C is a central component of the CCAN, responsible for interactions both with the CENP-A nucleosome on the chromatin-side as well as with subunits of the Mis12 complex on the kinetochore side [10]. Human CENP-C is a 934 amino acid long disordered protein, depletion of which leads to cell division defects and chromosome mis-segregation [11,12]. Two regions in human CENP-C have been identified as nucleosome binding regions: 1. the central region (aa 426-537), CENP-C^CR^, that is necessary and sufficient to promote CENP-A nucleosome binding *in vitro* and kinetochore targeting *in vivo* and 2. the CENP-C motif (aa 736-758), CENP-C^motif^, that is conserved across species, but is not sufficient for centromere targeting in the absence of endogenous CENP-C as it depends on the CENP-C dimerization domain. The CENP-C^motif^ is dispensable for epigenetic stability of the CENP-A nucleosomes [10,11,13].

The current molecular understanding of the CENP-A nucleosome/CENP-C interactions are based on the crystal structure of the canonical *D. melanogaster* nucleosome in which the C-terminal tail of histone H3 is replaced by the C-terminal tail of rat CENP-A, in complex with the rat CENP-C motif [14].

Here, we report a 3.8 Å cryoEM structure of the CENP-A nucleosome that confirms flexibility of DNA ends as an intrinsic property of CENP-A nucleosomes. We find that terminal DNA flexibility is independent of the nature of the underlining DNA sequence and is instead dictated primarily by the N-terminal tail of CENP-A. Furthermore, we find both nucleosome binding domains of CENP-C, CENP-C^CR^ and CENP-C^motif^, to be specific for CENP-A nucleosomes, where CENP-C^CR^ shows stronger binding. We also determined the cryoEM structure of the human CENP-A nucleosome in complex with human CENP-C^CR^ at 3.1 Å resolution and identified CENP-A^V532^ and CENP-A^V533^ as the key determinants for strong affinity of the CENP-A/CENP-C interaction. We notice conformational changes within the CENP-A nucleosome upon binding of CENP-C^CR^. The enhanced DNA unwrapping is facilitated by destabilization of the H2A C-terminal tail while the H4 N-terminal tail is stabilized in the conformation that favors centromere-specific H4^K20^ monomethylation.

In summary, our work provides a high-resolution, integrated view of the CENP-A nucleosome with its key CCAN partner, CENP-C. We establish CENP-A nucleosomes as the sole CENP-C binder and we provide a molecular understanding for the higher specificity of the CENP-C^CR^ compared to the CENP-C^motif^. Finally, our study identifies conformational changes in the nucleosome, taking place upon binding.

## Results

### 1. CENP-A nucleosome has flexible DNA ends, irrespective of DNA sequence

Ever since CENP-A has been identified as the key epigenetic mark of the centromere, a central question has been how it is distinguished from canonical nucleosomes [15]. Initial *in vitro* studies [5,6] together with recent research in cells, strongly support an octameric nucleosome, similar to the canonical one [16,17]. In the last 10 years, several studies both *in vivo* [16,18] and *in vitro* [6,18–20] have identified flexible DNA ends as a unique feature of CENP-A nucleosomes but to which degree DNA sequence and/or crystal packing contributed to unwrapping remained unclear. To determine this directly, we MNase digested CENP-A nucleosomes assembled both on synthetic super-positioning DNA “601” [21] and on two natural α-satellite DNA constructs [16], with and without the CENP-B box, a 17bp sequence recognized by CENP-B [22], respectively. Since the exact nucleosome positioning on the sequence that contains the CENP-B box is not precisely mapped, we used a full length α-satellite repeat (171 bp) with a CENP-B box at one of the ends. For all three DNA sequences, we observed faster DNA digestion when assembled on CENP-A nucleosomes in comparison to H3 nucleosomes (Figure 1A, Figure S1A). The DNA unwrapping of the CENP-A nucleosome has been linked to properties of the N-terminal sequence of CENP-A [18,23]. Indeed, when we substitute residues 1-49 of CENP-A with 1-50 of H3, we completely loose the DNA flexibility (Figure S1B). The converse is also true where H3 nucleosomes bearing the N-terminal tail of CENP-A undergo DNA unwrapping to a similar extent as CENP-A nucleosomes (Figure S1B). We therefore conclude that flexibility of the DNA ends is an intrinsic property of the CENP-A nucleosome that is regulated by its N-terminal tail and is independent of DNA sequence.

**Figure 1:**
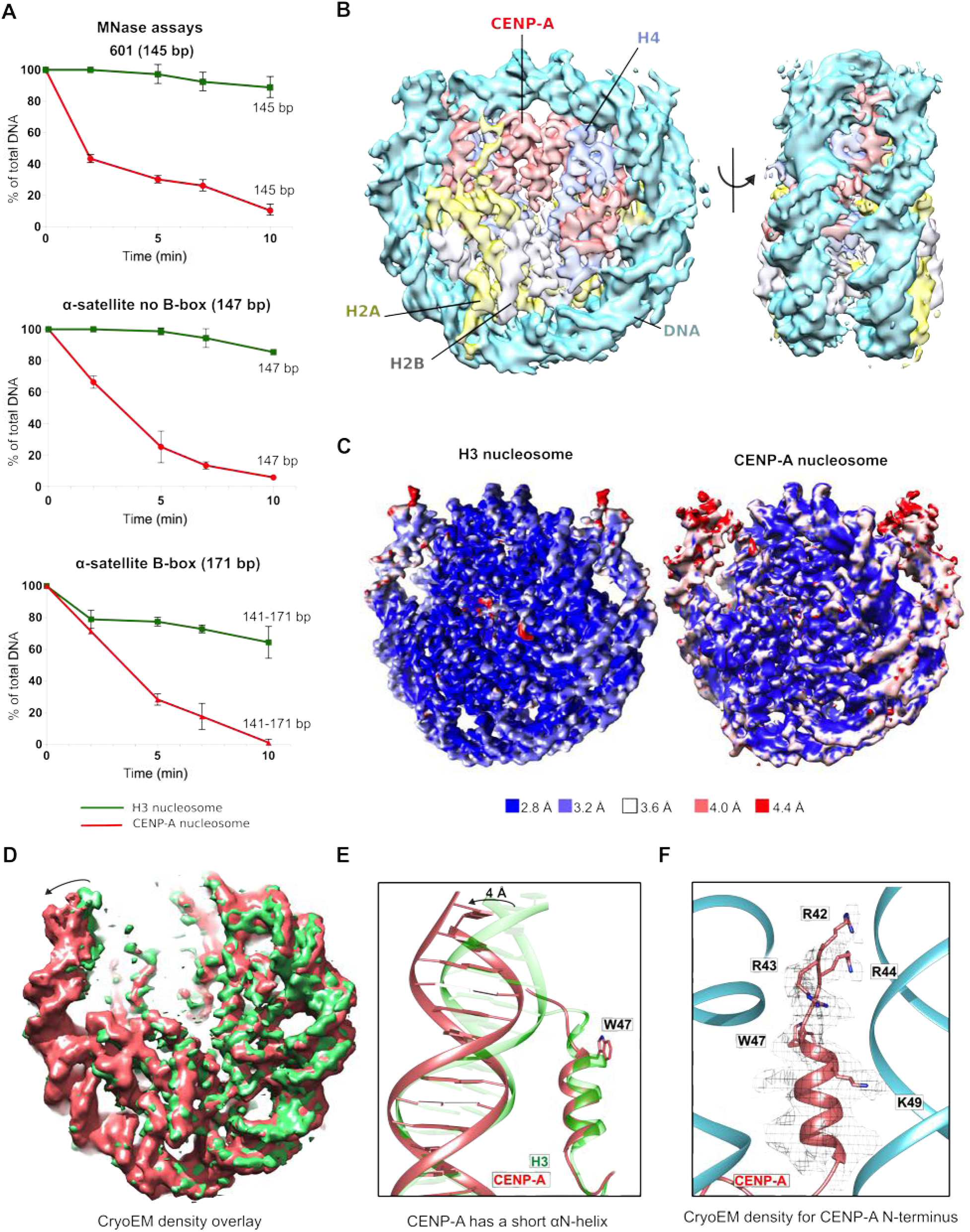
CENP-A nucleosome has flexible DNA ends independently of the wrapped DNA sequence. **A.** Graphs showing the relative abundance of undigested DNA as a function of time during digestion with micrococcal nuclease (MNase) for three types of DNA wrapped around the CENP-A nucleosome (red) and the H3 nucleosome (green). For α-satellite DNA with initial size of 171 bp, size ranges (141-171) corresponding to DNA lengths above NCP (nucleosome core particle) is presented. The standard error bar is shown for each time point based on three independent experiments. Virtual gels from Bioanaylzer are in Figure EV1A. **B.** CryoEM density map of the human CENP-A nucleosome, color-coded for histones and DNA. **C.** CryoEM maps of the H3 nucleosome (PDB 6ESF) and the CENP-A nucleosome, colored based on local resolution. **D.** Overlay of cryoEM maps of the H3 nucleosome (green; PDB 6ESF) and the CENP-A nucleosome (red). Note shorter and moved density for DNA on the CENP-A nucleosome indicated by the arrow. **E.** Overlay of the N-terminal tail of CENP-A (red) and H3 (green; PDB 6ESF), illustrating shorter αN helix of CENP-A (obstructed by the presence of bulky CENP-A^W47^) and DNA moved by 4 Å. **D.** The N-terminal tail of CENP-A (red) makes contacts with the DNA (cyan) at the entry/exit sites.

Next, we used cryoEM to obtain a high-resolution structure of CENP-A nucleosomes on 601 DNA at 3.8 Å (Figure 1B, Figure EV1C, Figure EV2, Table S1). In contrast to the crystal structure [6] where electron density for the terminal 13 DNA bp is missing, probably due to its flexible nature, our cryoEM structure of CENP-A nucleosomes reveals density for the entire 145 bp of DNA used in nucleosome reconstitutions. However, the map is less defined and has a lower local resolution for the terminal DNA (Figure 1C, Figure 1D and Figure SV2E), indicating local flexibility. The modeled DNA is shifted by 4 Å in comparison to the one in the H3 nucleosome [24] (Figure 1E). These results are in agreement with our MNase experiments and with a recent antibody-stabilized structure of the CENP-A nucleosome [20]. Interestingly, despite the low resolution for the DNA, we can clearly model the N-terminus of CENP-A all the way to CENP-A^R42^, including the αN helix (Figure 1F, Figure SV1D). The αN helix of CENP-A is shorter, disrupted by CENP-A^G46^ and the bulky CENP-A^W47^, while H3 continues with one extra turn (Figure 1E). CENP-A^R42^, CENP-A^R43^ and CENP-A^R44^ are all involved in DNA binding, although the interactions are slightly different on the two sides of the nucleosome (Figure 1F, Figure EV1D). In our structure, we can also clearly see other CENP-A specific features of the nucleosome (Figure EV1D): The C-terminal tail (-LEEGLG), that specifically binds CENP-C and L1 loop containing CENP-A^R80^ (“RG-loop”), that specifically binds CENP-N.

### 2. CENP-C^CR^ competes out the CENP-C^motif^ on CENP-A nucleosomes

Initial efforts to identify CCAN proteins that directly and specifically bind CENP-A nucleosomes uncovered the CENP-C and CENP-N proteins [4,25]. In those studies, only the central region of human CENP-C (CENP-C^CR^, aa 426-537) was characterized for CENP-A binding (Figure 2A). In 2013, Kato at al. [14] reported a crystal structure of the canonical H3 nucleosome in which the C-terminal tail of H3 was replaced by a mutated rat CENP-A C-terminus in complex with the rat CENP-C residues 710-734, a region known as the CENP-C motif (CENP-C^motif^). Analysis of the structure identified the hydrophobic interactions between the CENP-A C-terminal tail and the CENP-C^motif^, and the authors proposed that both CENP-C^CR^ and CENP-C^motif^ bind nucleosomes, with 5-10 times higher affinity for CENP-A then for H3 nucleosomes. To determine if CENP-C^CR^ binds the nucleosome with a different affinity than the CENP-C^motif^, we prepared complexes of both and analyzed their mobility on native PAGE. We found that both the CENP-C^CR^ and the CENP-C^motif^ complexes with CENP-A nucleosomes in a ∼2:1 ratio and travel as a sharp band on the native gel, corresponding to a uniform complex (Figure 2B, 2C). Surprisingly, when mixed with H3 nucleosomes, both CENP-C constructs result in a smear on the native gel, indicating non-specific binding (Figure 2B, 2C). We conclude that *in vitro* CENP-C^CR^ and CENP-C^motif^ bind only CENP-A nucleosomes specifically.

**Figure 2:**
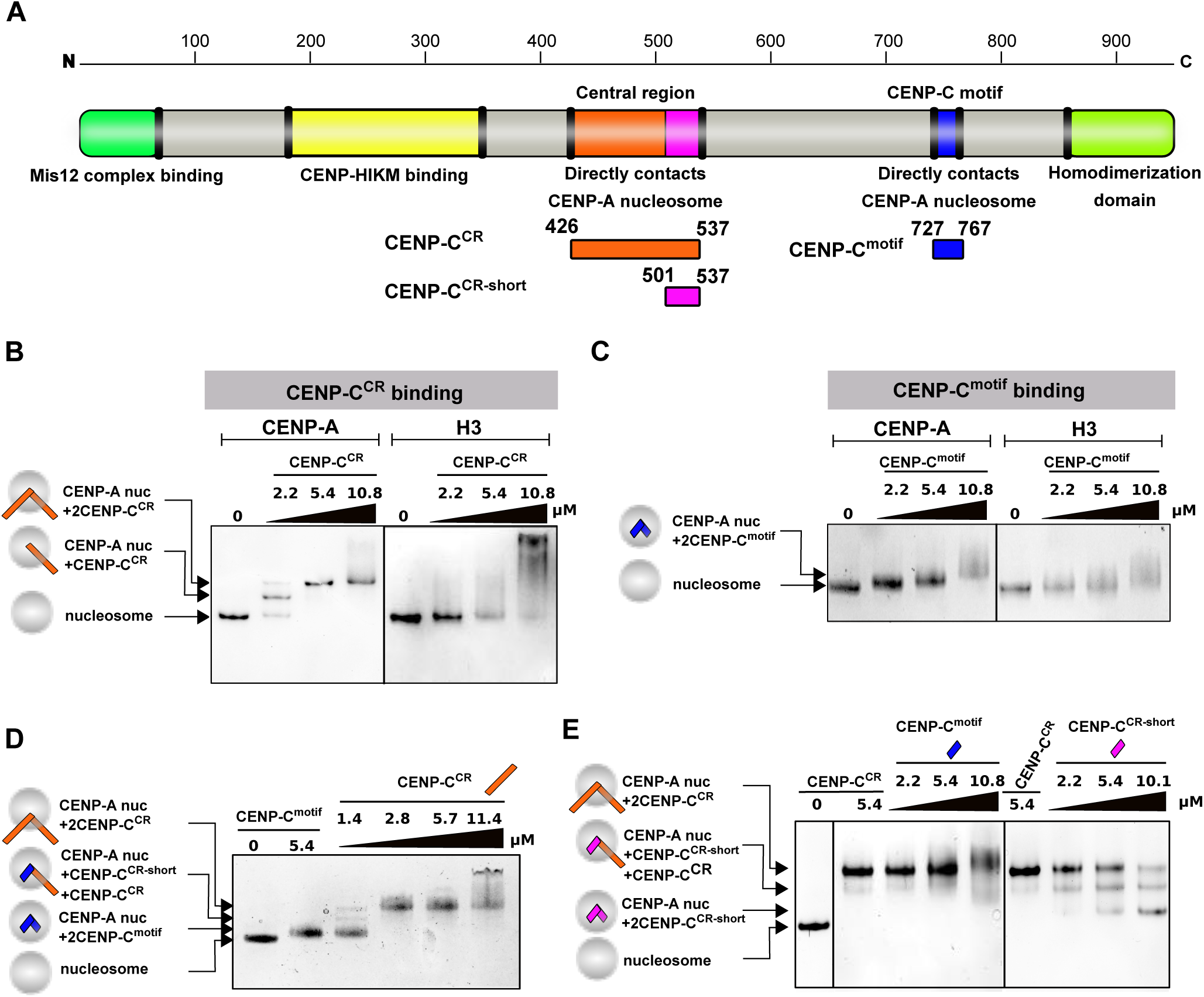
Both CENP-C^CR^ and CENP-C^motif^ bind specifically to the CENP-A nucleosome and CENP-C^CR^ easily competes out CENP-C^motif^ bound to CENP-A. **A.** Schematic diagram of the full-length CENP-C protein, indicating parts involved in interactions with other proteins or homo-dimerization. Constructs used in this study are depicted below the diagram. **B.** Native PAGE gel stained with EtBr showing complexes formed between CENP-A or H3 nucleosome and CENP-C^CR^. Lane 1: CENP-A nucleosome, Lanes 3-5: Increasing amounts of CENP-C^CR^ are added to CENP-A nucleosome. Generation of a sharp band with slower mobility indicates formation of a specific CENP-A/CENP-C^CR^ complex. Lane 5: H3 nucleosome. Lanes 6-8: Increasing amounts of CENP-C^CR^ are added to H3 nucleosome. Smear on the gel indicates formation of non-specific CENP-A/CENP-C^CR^ complexes. **C.** Same experiment as in B using CENP-C^motif^. **D.** Native gel showing CENP-C^CR^ competing out CENP-C^motif^ bound to CENP-A nucleosome. Lane 1: CENP-A nucleosome, Lane 2: CENP-A/CENP-C^motif^ complex. Lane 3-6: Increasing amounts of CENP-C^CR^ are added to the pre-formed CENP-A/CENP-C^motif^ complex. Formation of slower migrating bands indicates that longer CENP-C^CR^ is replacing shorter CENP-C^motif^ bound to the CENP-A nucleosome. **E.** Native gel showing the inability of CENP-C^motif^ to compete out CENP-C^CR^ bound to CENP-A nucleosome. Lane 1: CENP-A nucleosome. Lane 2 and 6: CENP-C^motif^/CENP-A nucleosome complex. Lanes 3-5: Increasing amounts of CENP-C^motif^ are added to the pre-formed CENP-A/CENP-C^CR^ complex. Formation of smear at high amounts of CENP-C^motif^ added, indicates that CENP-C^motif^, at high concentrations, non-specifically binds CENP-A/CENP-C^CR^ complex rather than replacing bound CENP-C^CR^. Lane 7-9: Increasing amounts of CENP-C^CR-short^ are added to the pre-formed CENP-A/CENP-C^CR^ complex. Formation of bands with higher mobility indicates that smaller CENP-C^CR-short^ is effectively replacing bigger CENP-C^CR^ bound to CENP-A nucleosome. For B-E, 2.4 μM nucleosomes are used in all experiments.

Furthermore, we performed a competition experiment to determine which CENP-C region has higher affinity for CENP-A nucleosomes. When we preassemble the human CENP-A/CENP-C^motif^ complex and titrate in CENP-C^CR^, we observe formation of the CENP-A/CENP-C^CR^ complex with almost full saturation at 2 x molar access of CENP-C^CR^ (Figure 2D). In contrast, titrating in the CENP-C^motif^ to a pre-formed CENP-A/CENP-C^CR^ complex, results in a smear on the native PAGE, indicative of non-specific binding of CENP-C^motif^ to CENP-A/CENP-C^CR^ complexes (Figure 2E). The CENP-C^motif^ is 71 residues shorter than CENP-C^CR^ (40 vs. 111 residues) and it is possible that extra residues, beyond those directly interacting with the CENP-A C-terminus are providing additional affinity. To test this, we made a truncated CENP-C^CR^ that has only residues predicted to interact with the CENP-A C-terminal tail (501-537; CENP-C^CR-short^) and is comparable in size to the CENP-A^motif^. First, we confirmed specificity of CENP-C^CR-short^ for CENP-A nucleosomes (Figure EV3). Interestingly, we also observed that CENP-C^CR-short^ completely loses its ability to bind H3 nucleosomes. This suggests that the additional 71 residues present in the CENP-C^CR^ are responsible for the non-specific H3 nucleosome binding, likely through interactions with DNA. Next, we tested if CENP-C^CR-short^ can compete out CENP-C^CR^ from pre-assembled CENP-A/CENP-C^CR^ complexes. We find that, in contrast to the CENP-C^motif^, CENP-C^CR-short^ efficiently replaces CENP-C^CR^ bound to CENP-A nucleosomes (Figure 2E), demonstrating that residues within the nucleosome binding region of CENP-C^CR^ contribute to high affinity binding.

In summary, we conclude that CENP-C binds only CENP-A nucleosomes, and that CENP-C^CR^ provide the major interactions between CENP-A and CENP-C. Our findings are consistent with studies in cells that found CENP-C^CR^ to be essential for epigenetic propagation of the centromere [13].

### 3. Strong hydrophobic interactions are contributing specificity and high affinity between the CENP-A nucleosome and CENP-C^CR^

Next, we aimed to define how the high affinity CENP-C^CR^ interaction is achieved. The crystal structure of the H3-GIEGGL/rat CENP-C^motif^ has identified two types of interactions involved in complex formation: 1. electrostatic interactions of rat CENP-C^R717/R719^ with the acidic patch on H2A/H2B, and 2. hydrophobic interactions between rat CENP-C^Y725, W726^ and the hydrophobic C-terminal tail of CENP-A [14]. A sequence comparison of the rat CENP-C^motif^ with the human CENP-C^motif^ and human CENP-C^CR^ reveals the existence of analogous residues in the human protein but does not explain the higher affinity of the CENP-C^CR^ (Figure 3A). Also, since crystallographic studies [14] used a canonical nucleosome with a grafted C-terminus of CENP-A, it failed to capture possible conformational changes that could occur within the CENP-A nucleosome upon CENP-C binding. To resolve this, we solved the 3.1 Å cryoEM structure of the human CENP-A nucleosome in complex with human CENP-C^CR^ (Figure 3B, Table S2, Figure EV4, Figure EV5). Our maps show CENP-C bound to both sides of the nucleosome via the CENP-A^R521, R522^ anchoring residues, while the remaining residues in CENP-C^CR^ show fuzzy interactions. In order to visualize a larger CENP-C fragment, we have further sorted particles and increased CENP-C occupancy on one of the sides, so that we can trace residues 519-536. We observe strong electrostatic interactions between human CENP-C^R521, R522^ and the acidic patch formed by H2A/H2B (Figure 3C, Figure EV4A). Neutralization of a negative surface contributed by H2A/H2B on the nucleosome is a feature common to a handful of other nucleosome binding molecules (reviewed in [26]) that was also seen in the chimeric H3 nucleosome/CENP-C^motif^ structure [14]. We also observe human CENP-C^W530, W531^ tightly fitting the hydrophobic cleft formed by the C-terminal tail of CENP-A (Figure 3C, Figure EV4B). Furthermore, the conformation of the C-terminal tail of CENP-A changes slightly upon binding (Figure 3D), and we observe more extended hydrophobic interactions in comparison to those reported in the crystal structure (Figure 3E). Two bulky tryptophans (CENP-C^W530, W531^) are clamped between CENP-A^R131^ and CENP-A^L135, L139^ while CENP-C^V532^ and CENP-C^V533^ stabilize the hydrophobic patch on H4 formed by H4^V58^ and H4^V61^. These interactions explain the perturbations observed by NMR in H4 residues within the nucleosome upon CENP-C^CR^ binding [14]. To test if extended hydrophobicity contributed by CENP-C^V532, V533^ plays a role in stronger CENP-A nucleosome/CENP-C^CR^ interactions, we generated a CENP-C mutant devoid of residues 530-537, CENP-C^CR-ΔC^ (Figure 3F). This mutant fails to form a complex with the CENP-A nucleosome (Figure 3G), indicating that the residues following CENP-C^W530, W531^ are important for binding. Point mutations further revealed that replacement of CENP-C^V533^ with aspartic acid is sufficient to eliminate binding to the CENP-A nucleosome (Figure 3F, Figure 3G).

**Figure 3.**
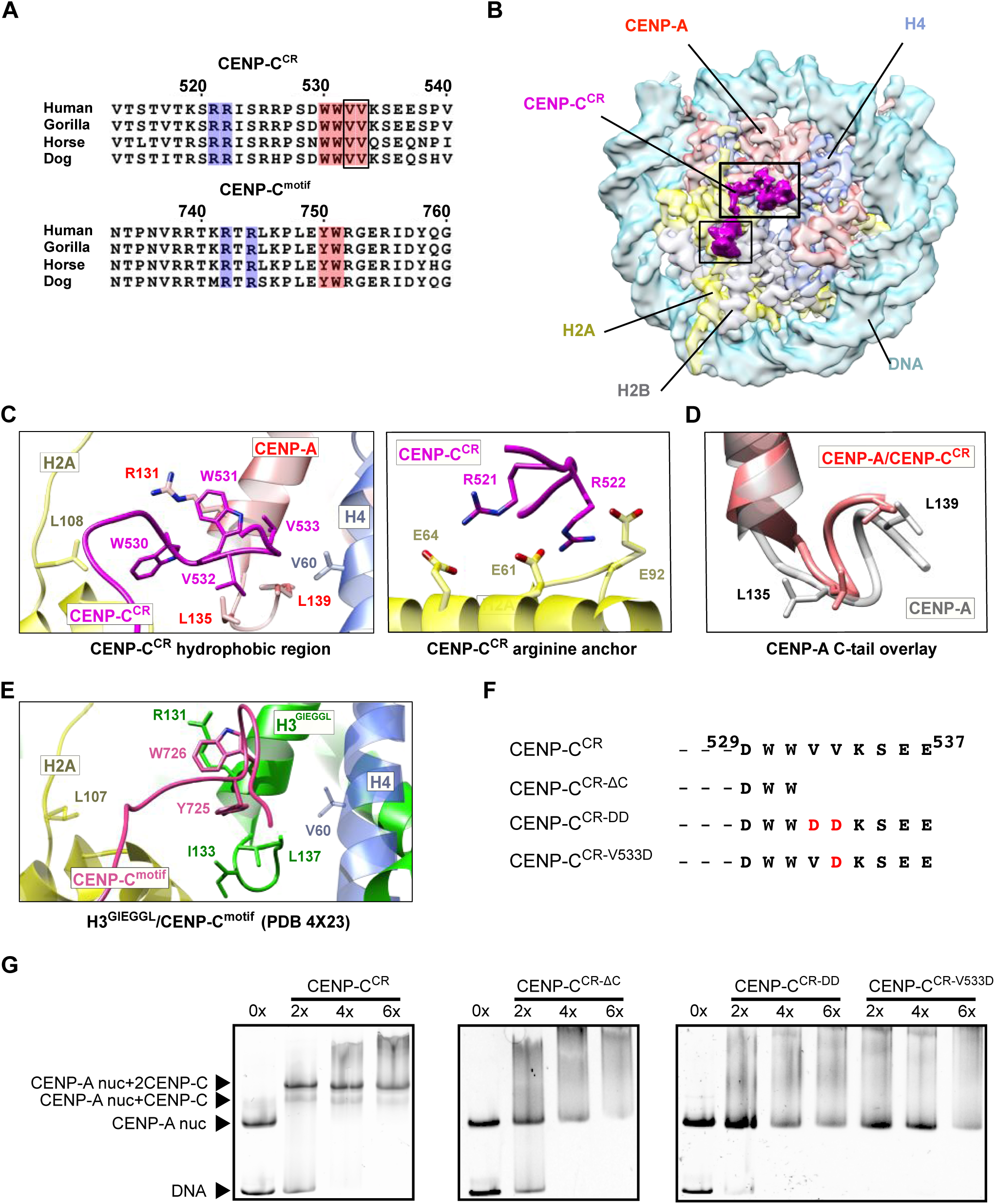
Strong hydrophobic interactions are contributing specificity and high affinity between the CENP-A nucleosome and CENP-C^CR^. **A.** Sequence alignment of CENP-C^CR^ and CENP-C^motif^ regions from different mammals. Conserved residues involved in CENP-A binding are highlighted in blue (electrostatic interactions) and pink (hydrophobic interactions). Residues identified in this study to contribute higher affinity of CENP-C^CR^ comparing to CENP-C^motif^, are boxed. **B.** CryoEM density map of the human CENP-A nucleosome, color-coded for histones, in complex with CENP-C^CR^ (magenta). **C.** Ribbon diagram showing interactions of the CENP-C^CR^ hydrophobic region (left) and CENP-C^CR^ arginine anchor (right) with the CENP-A nucleosome. **D.** Overlay of the CENP-A C-terminal tail before (gray) and after (red) binding of CENP-C^CR^. **E.** Interactions between H3-GIEGGL and rat CENP-C^motif^ as observed in PDB 4X23. **F.** Schematic diagram of mutated CENP-C^CR^ sequences used to test importance of CENP-C^V532^ and CENP-C^V533^ for generation of CENP-A/CENP-C^CR^ complexes. **G.** Native gels showing the impaired ability of mutated CENP-C^CR^ to form complexes with the CENP-A nucleosome. The molar ratio of CENP-C^CR^/CENP-A nucleosomes is shown above each lane.

We conclude that human CENP-C^CR^ binds the human CENP-A nucleosome through electrostatic interactions, involving CENP-C^R521, R522^ (neutralizing the acidic patch on the nucleosome) as well as via an extensive hydrophobic network formed between CENP-C^W530, W531, V532, V533^ and the C-terminal tail of CENP-A, aided by the hydrophobic patch of H4. Our data show that CENP-C^CR^ makes additional hydrophobic contacts with the CENP-A nucleosome that have not been observed for CENP-C^motif^. These interactions are required for the higher affinity of CENP-C^CR^ versus the CENP-C^motif^.

### 4. Conformational changes within the CENP-A nucleosome upon CENP-C binding

The nucleosome in our experiments bears the full length CENP-A, which enables us to assess conformational changes upon CENP-C binding. It has previously been proposed that CENP-C binding rigidifies the histone core, rendering it more histone H3-like, while at the same time further enhancing DNA unwrapping [27]. The MNase experiments on CENP-A nucleosomes in complex with CENP-C^CR^ or CENP-C^motif^ show increased DNA digestion in both cases (Figure 4A, Figure EV6A, left). Comparison of the local resolution maps between CENP-A and CENP-A/CENP-C^CR^ complex (Figure S3E; Figure S5E) indicates more extensive DNA unwrapping in samples with bound CENP-C^CR^. To further confirm this, we performed careful cryoEM classification of particles based on the conformations at DNA entry/exit sites for each of the two samples (Figure S3E; Figure S5E). We find that, although flexible, the DNA at the entry/exit site in CENP-A nucleosome samples only a limited space (distance between most extreme DNA conformation is 2 Å) (Figure 4B). In contrast, in the CENP-A/CENP-C^CR^ complex the most extreme distance between different subpopulations is 9 Å (Figure 4C). Concomitantly, we observe no density for the C-terminal tail of H2A (Figure 4C), indicating disorder in this part of the nucleosome upon CENP-C binding. This is in striking contrast to the cryoEM structure of the canonical nucleosome with tightly wrapped DNA, where the density for the C-terminal tail of H2A is one of the most well resolved parts of the structure [24]. It has been reported that removal of the C-terminal tail of H2A leads to decreased stability of the nucleosome, alters nucleosome positioning and interactions with H1 [28,29]. Molecular dynamic studies have identified residues H2A^K118^ and H2A^K119^ to interact with DNA, most likely securing the DNA wrap [30]. Indeed, if we remove residues 110-130 in H2A from the CENP-A nucleosome (CENP-A^ΔC-tail H2A^), we observe increased DNA digestion (Figure 4D, Figure EV6B, top), and addition of CENP-C^CR^ to these nucleosomes does not change the nuclease digestion profile (Figure 4D, Figure EV6B, bottom). In contrast, DNA digestion of H3 nucleosomes was only mildly affected by removal of the C-terminus of H2A (Figure EV6C, top). We next hypothesized that the different MNase pattern of CENP-A vs. H3 nucleosomes (in the background of ΔC-tail H2A) must be contributed by the residues in the N-terminal tail of CENP-A/H3, since this region of the nucleosome is in close contact with the C-terminal tail of H2A. Indeed, in the context of ΔC-tail H2A, MNase digestion of nucleosomes with an H3 core and CENP-A tail is highly similar to CENP-A nucleosomes while digestion of nucleosomes with a CENP-A core and H3 tail is similar to that of the H3 nucleosome (Figure EV6C, middle). These results demonstrate a contribution of the C-terminal tail of H2A to nucleosome DNA wrapping that synergizes with the N-terminal tail of CENP-A. In the context of the CENP-A nucleosome that has a shorter αN helix, DNA wrapping is already compromised and removal of C-terminal H2A results in further unwrapping (Figure 4E). Consistently, we find that in CENP-A nucleosomes with an H3 N-tail, binding of CENP-C^CR^ alone cannot induce DNA unwrapping (Figure EV6C, bottom). From this, we conclude that the N-terminal tail of CENP-A favors DNA unwrapping which is counteracted by the C-terminus of H2A. CENP-C binding destabilizes the C-terminus of H2A, leading to increased DNA unwinding. We find that this alternative, destabilized, conformation of the H2A C-terminal tail is most likely caused by an interaction between the bulky hydrophobic CENP-C^W530, W531^ and H2A^L108^. In addition, CENP-C^W530, W531^ is orienting CENP-A^R131^ to establish a salt bridge with H2A^Q112^ (Figure EV6D).

**Figure 4.**
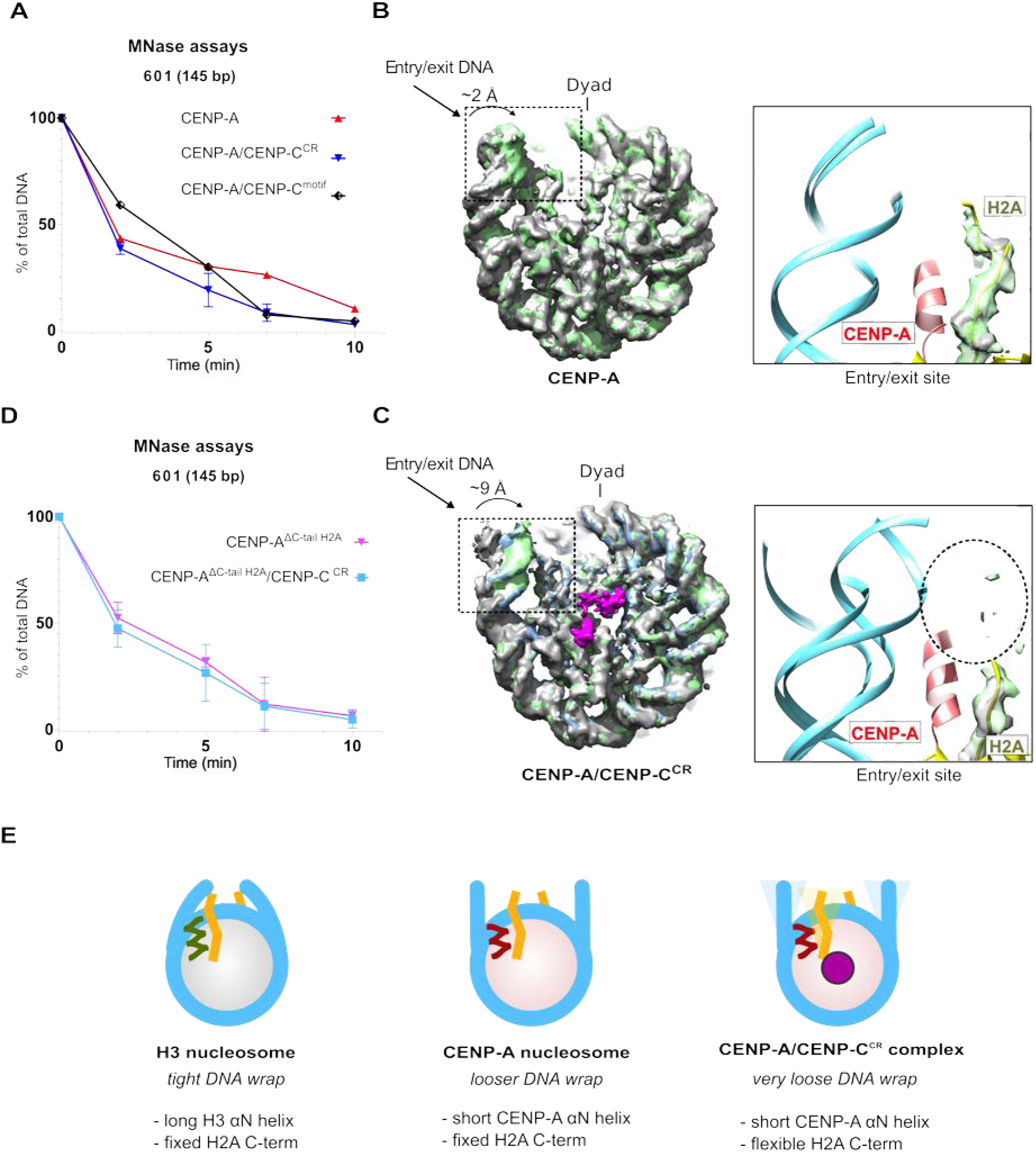
The C-terminus of H2A is destabilized in the CENP-A/CENP-C^CR^ complex. **A.** Graph showing the relative abundance of undigested DNA (145 bp) as a function of time during digestion with micrococcal nuclease (MN-ase) for the CENP-A nucleosome (red), the CENP-A/CENP-C^CR^ complex (blue) and the CENP-A/CENP-C^motif^ complex (black). **B.** Overlay of three cryoEM maps of the CENP-A nucleosome obtained by sorting on the DNA entry/exit site (Figure EV2F). Distance between most open (grey) and most closed (green) map is 2 Å. The entry/exit site is boxed and the corresponding model is shown on the right. Note that the density of the H2A C-terminus is well defined. **C.** Overlay of three cryoEM maps of the CENP-A/CENP-C^CR^ complex obtained by sorting on the DNA entry/exit site (Figure EV5F). The distance between the most open (grey) and most closed (green) maps is 9 Å. The entry/exit site is boxed and the corresponding model is shown on the right. Note the absence of a clearly defined density of the H2A C-tail. **D.** Same type of data as in A. for the CENP-A nucleosome assembled with tailless H2A (H2A, 1-109) alone (pink) and in complex with CENP-C^CR^ (light blue). **E.** Schematic representation of the interplay between the N-terminus of H3 or CENP-A and the C-terminus of H2A in regulating flexibility of nucleosomal DNA wrap. DNA (cyan); longer H3 αN (green); shorter CENP-A αN (red); H2A C-terminus (yellow); CENP-C (magenta). For A and C. standard error bars are shown for each time point based on three independent experiments and virtual gels are available in Figure EV6.

Furthermore, we noticed that the N-terminal tail of H4 clearly adopts an upwards conformation on both sides of the CENP-A nucleosome relative to canonical nucleosomes (Figure 4A, left), but in the presence of CENP-C^CR^ this conformation is more rigidified (Figure 4A, right). Monomethylation of H4 is enriched on CENP-A nucleosomes and it is necessary for epigenetic establishment of the kinetochore [31]. A recent structural study [32] proposed an upward conformation of the H4 tail to be required for establishment and/or maintenance of H4^K20^ monomethylation. In our cryoEM maps, we observe the N-terminus of H4 in a slightly different but still upwards conformation in comparison to the crystal structure of the H3^CATD^ nucleosome [32]. A very recent study [33], indicates that the conformation of the N-terminus of H4 is further modified upon CENP-N binding.

In summary, we find that binding of CENP-C^CR^ to the CENP-A nucleosome enhances DNA unwrapping by destabilizing the C-terminal tail of H2A while, at the same time, stabilizing the H4 C-terminal tail in the upward conformation that may be important for centromere specific monomethylation of H4^K20^.

## 5. Discussion

CENP-A is a key epigenetic mark to maintain centromere identity, a chromosome locus essential for genome integrity. CENP-A is a histone H3 variant with a histone core bearing 64% identity to H3 while featuring completely divergent tails. As CENP-A is a central epigenetic determinant of the centromere, an important question is how CENP-A containing nucleosomes are distinguished from bulk chromatin. Over the years, a number of models have been proposed, including different histone stoichiometries, presence of non-histone proteins and alternative DNA wrapping [summarized in [15]]. Finally, an octameric nucleosome with a right-handed DNA wrap, much like canonical nucleosomes, emerged as the favorite model for the human CENP-A nucleosome, based on both *in vitro* and *in vivo* studies [5,6,16,17], leaving open the question of what is making CENP-A nucleosomes so unique. A crystal structure of the CENP-A nucleosome [6], *in vivo* MNase experiments [16] and initial cryoEM studies [18] all pointed towards enhanced DNA unwrapping as a CENP-A specific feature where unwrapped DNA results in a different chromatin architecture, potentially important for accommodating CCAN components. Indeed, the experiments presented here, together with the structure of CENP-A, confirm high flexibility of the DNA ends as an intrinsic feature of CENP-A nucleosomes encoded in the N-terminal tail of CENP-A, independent on the DNA sequence.

A key question is how the CENP-A nucleosome directly binds two CCAN components, CENP-N and CENP-C, and how their binding is changing the nucleosome. Recent work [7–9] has provided a view of the CENP-A/CENP-N interaction at atomic resolution, which involves recognition of the solvent-exposed, positively charged L1 loop on CENP-A and an interaction with DNA, with minimal changes to the rest of the nucleosome. For the CENP-C interactions with the CENP-A nucleosome, 2 different parts of CENP-C are proposed to bind: CENP-C^CR^ and CENP-C^motif^. CENP-C^CR^ is necessary and sufficient for CENP-A nucleosome binding *in vitro* and centromere targeting and stability *in vivo* while the CENP-C^motif^ can be recruited to kinetochores only in the presence of a homo-dimerization domain [10,11,13]. Having two independent CENP-C modules able to bind nucleosome at centromeres with different affinities, led to the proposal [10] of CENP-C, as a direct mediator of CENP-A loading during the cell cycle. The model assumes that the CENP-C module with weaker nucleosome binding, CENP-C^motif^, alternates between binding of an H3.3 nucleosome and a CENP-A nucleosome while the stronger module, CENP-C^CR^, remains stably associated with the CENP-A nucleosome in all phases of the cell cycle. We show that, indeed, CENP-C^CR^ binds the CENP-A nucleosome with higher affinity than CENP-C^motif^, but neither of the CENP-C modules can make uniform complexes with H3 nucleosome.

Furthermore, structural insights of the CENP-A/CENP-C interactions are based on the crystal structure of a chimeric fruit fly H3 nucleosome with the C-terminal tail of rat CENP-A and interaction of this tail with the rat CENP-C^motif^ [14]. We here report a 3.1 Å structure of a complete human CENP-A nucleosome in complex with human CENP-C^CR^ which is essential for epigenetic stability of centromeres [13]. We find a longer hydrophobic stretch on CENP-C^CR^, formed by CENP-C^V532, V533^, to be essential for robustness of the CENP-A nucleosome/CENP-C^CR^ interactions, and we re-define the minimal fragment of CENP-C^CR^ necessary for productive CENP-A binding (residues 501-537).

Previous experiments [27] and those reported here, confirm enhanced DNA unwrapping of the CENP-A nucleosome, induced by CENP-C^CR^ binding. In our structure of the CENP-A/CENP-C^CR^ complex, we observe a disordered H2A C-terminal tail and using a combination of MNase experiments and mutagenesis we establish a role for the H2A histone tail in regulating the extent of DNA wrapping on nucleosomes. We find that the C-terminal tail of H2A secures nucleosomal DNA wrapping that counteracts the unwrapping of CENP-A nucleosomes promoted by the short CENP-A αN helix. Upon CENP-C binding the hydrophobic interaction between H2A and CENP-C is displacing the H2A C-terminal tail, resulting in a very loose DNA wrapping in the CENP-A/CENP-C^CR^ complex. Regulation of the DNA wrap by the C-terminal tail of H2A is likely exploited in general chromatin. For example, H2A variants with different C-terminal tails are known to regulate various biological processes, conferring special properties to the chromatin [34].

Furthermore, in both our structures of the CENP-A nucleosome alone and the CENP-A/CENP-C^CR^ complex, we see an upwards conformation of the C-terminal tail of H4. The conformation is additionally stabilized by CENP-C^CR^ binding, most likely through hydrophobic interactions between CENP-C and H4. H4^K20^ monomethylation is essential for kinetochore assembly [31,32], and binding of CENP-C^CR^ could be enforcing this centromere-specific epigenetic chromatin modification.

Combined, our structures provide an essential and long anticipated high-resolution view of the fundamental building block of the centromere, the CENP-A nucleosome alone and in complex with CENP-C, a protein that forms the backbone of the constitute centromere complex. Our biochemical and structural analysis establishes CENP-C as an exclusive and multivalent binder of CENP-A nucleosomes that employs two independent modules and homo-dimerization to crosslink sparse CENP-A domains in centromeric chromatin [35,36] and provide framework for functional CCAN (Figure 6). Binding of CENP-C to CENP-A nucleosomes, not only spatially organizes CENP-A nucleosomes and recruits other CCAN components, it also induces conformational changes (DNA unwrapping, neutralization of the acidic patch on H2A and facilitation of H4^K20^ monomenthyation) that might be required for establishment and maintenance of functional centromeres (Figure 6).

**Figure 5.**
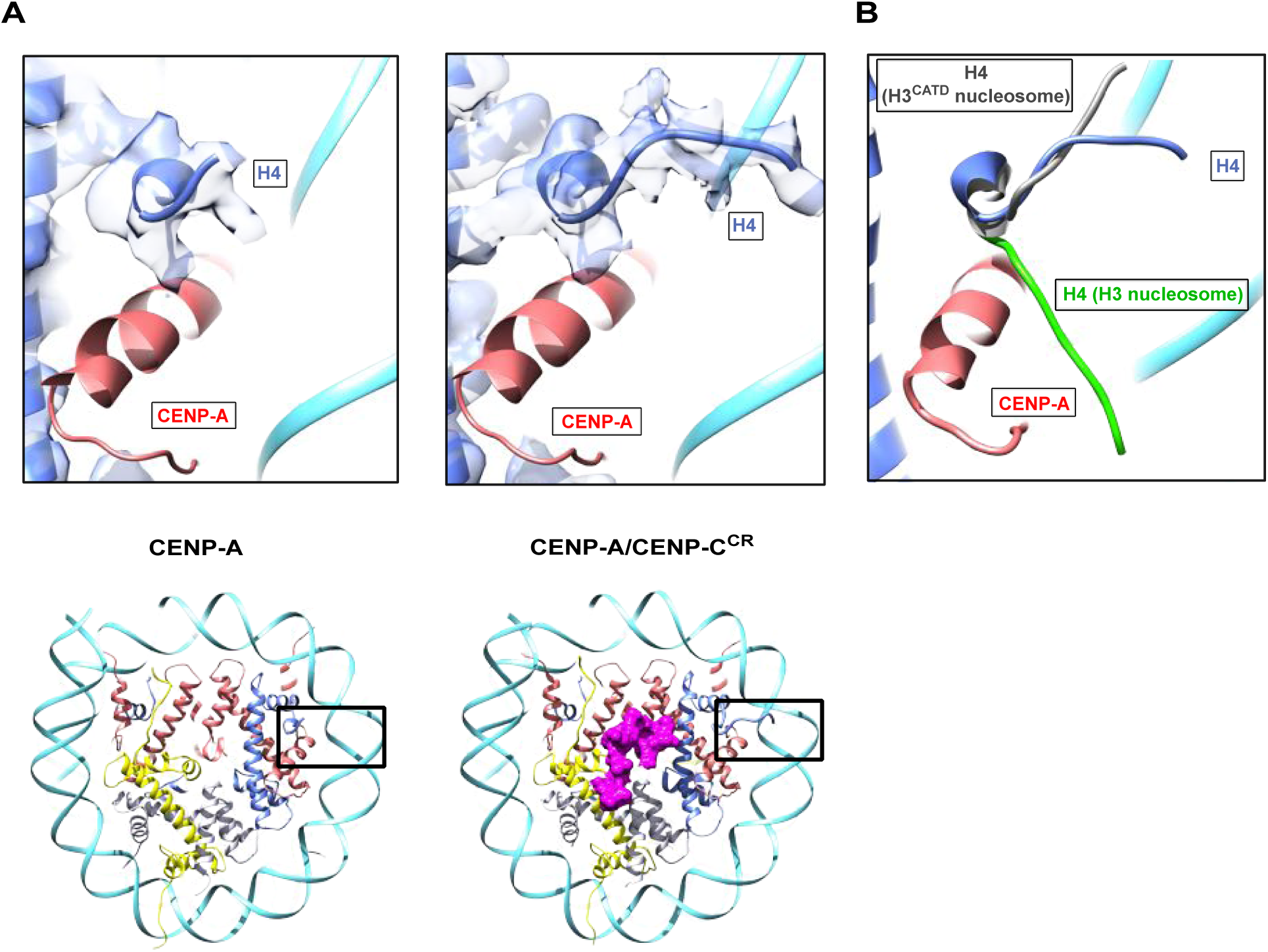
The N-terminal tail of H4 is stabilized in upwards conformation in the CENP-A/CENP-C^CR^ complex. **A.** Structure of the H4 N-terminal tail with corresponding density in the CENP-A nucleosome (right) and CENP-A/CENP-C^CR^ complex (left). **B.** Overlay of the H4 N-tail from the CENP-A/CENP-C^CR^ complex (blue) with the H4 N-terminus from the H3 nucleosome (green; PDB 6ESF) and H3^CATD^ (grey; PDB 5Z23).

**Figure 6.**
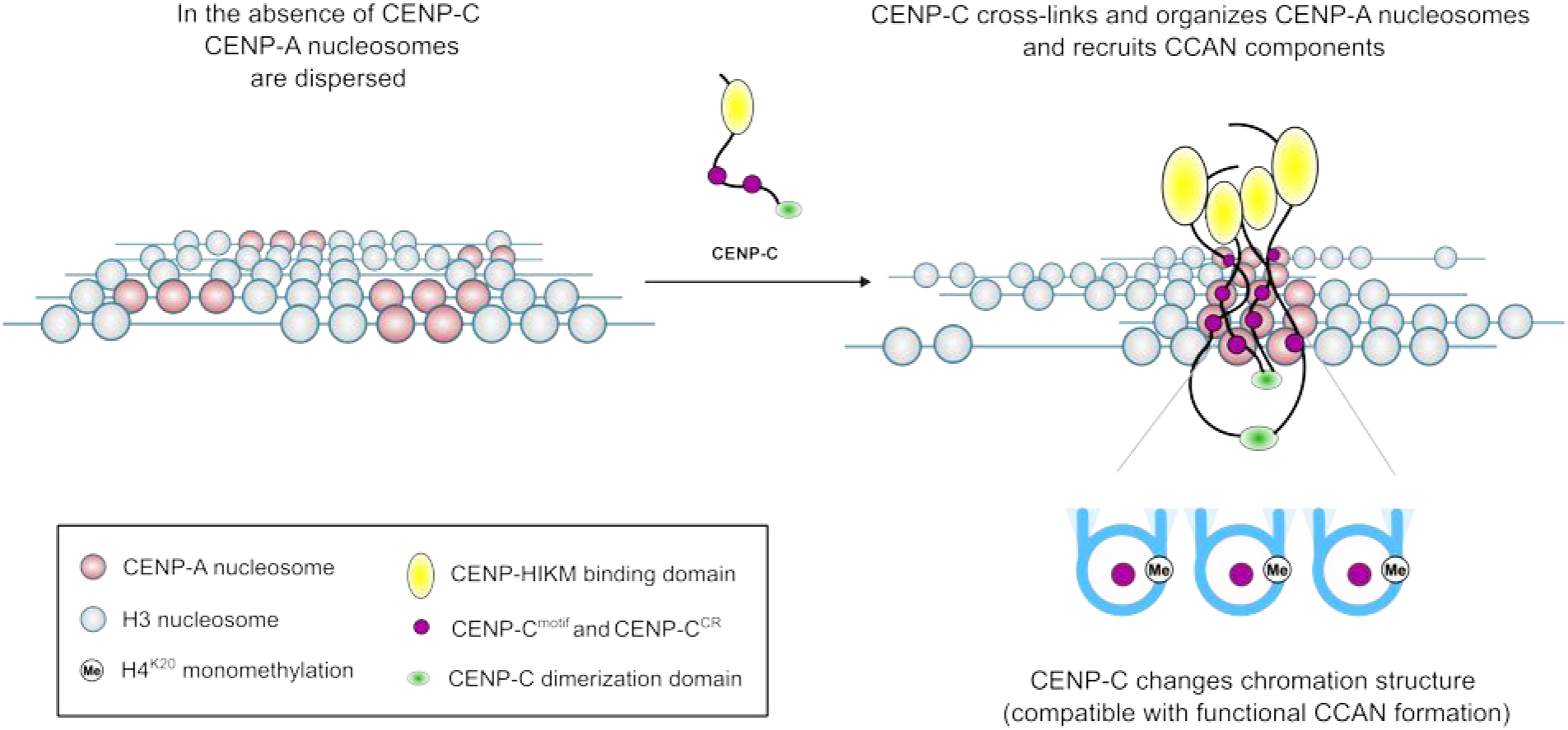
Model illustrating role of CENP-C in centromeric chromatin. CENP-C crosslinks CENP-A nucleosomes, recruits other CCAN components and pre-conditions chromatin for formation of a functional centromere.

## Materials and methods

### Protein purification

Human histones, CENP-A and the CENP-C central domain, were expressed and purified as previously described in [37]. Briefly, the CENP-A/H4 hetero-tetramer was expressed from a bicistronic plasmid in *E. coli* pLysS under soluble conditions and purified on a hydroxyapatite column followed by cation exchange chromatography. H3, H4, H2A and H2B were expressed in inclusion bodies and purified under denaturing conditions using a Sephacryl size exclusion column followed by cation exchange chromatography. H2A/H2B and H3/H4 histones were subsequently co-refolded to form hetero-dimers and hetero-tetramers, respectively, and purified with size exclusion chromatography.

GST-tagged recombinant human CENP-C central region (CENP-C^CR^, aa 426–537), CENP-C motif (CENP-C^motif^, aa 727-767) and short CENP-C central region (CENP-C^CR-short^, aa 501-537) were expressed and affinity-purified on a glutathione column. GST was subsequently cleaved overnight by PreScission protease and separated from CENP-C, using cation exchange and size exclusion chromatography.

PCR site-directed mutagenesis was performed to generate CENP-C^V532D/V533D^, CENP-C^V533D^, CENP-C^426-531^ and CENP-C^426-533^. All mutants were expressed and purified as described for the central CENP-C region.

The H3 histone with a CENP-A N-tail^1-49^ (H3^CENP-A (N-tail)^), CENP-A histone with a H3 N-tail^1-50^ (CENP-A^H3(N-tail)^) (figure S1B) and a H2A histone lacking the C-terminal residues 110-130 were cloned using In-Fusion® HD cloning strategy (Takara Bio).

145 bp 601 super-positioning DNA and 147 bp α-satellite DNA were purified as described in [38]. Briefly, XL10 cells transformed with pUC57 plasmids containing 6 × 147 bp α-satellite DNA and 8 × 145 bp 601 super-positioning DNA (gift from Ben Black, UPenn) were cultivated, DNA was extracted with phenol/chloroform, digested using EcoRV restriction enzyme and further purified using anion-exchange chromatography. 171 bp α-satellite DNA with a CENP-B box was amplified from plasmid using PCR and further purified using anion-exchange chromatography.

601 (145 bp):

ATCAGAATCCCGGTGCCGAGGCCGCTCAATTGGTCGTAGACAGCTCTAGCACCGCTTAAACGCACGTACGCGCTGTCCCCCGCGTTTTAACCGCCAAGGGGATTACTCCCTAGTCTCCAGGCACGTGTCAGATATATACATCGAT

α-satellite no B-box (147 bp):

ATCAAATATCCACCTGCAGATTCTACCAAAAGTGTATTTGGAAACTGCTCCATCAAAAGGCATGTTCAGCTCTGTGAGTGAAACTCCATCATCACAAAGAATATTCTGAGAATGCTTCCGTTTGCCTTTTATATGAACTTCCTCGAT

α-satellite **<u>B-box</u>** (171 bp):

GGAGGA**<u>TTTCGTTGGAAACGGGA</u>**TCAACTTCCCATAACTGAACGGAAGCAAACTCAGAACATTCT TTGTGATGTTTGTATTCAACTCACAGAGTTGAACCTTCCTTTGATAGTTCAGGTTTGCAACACCCTTGTAGTAGAATCTGCAAGTGTATATTTTGACCACTTTGGA

### Assembly of nucleosomes and nucleosome complexes

CENP-A and H3 nucleosomes were assembled using 601 (145 bp), α-satellite no B-box (147 bp) or α-satellite with B-box (171 bp) DNA. H2A/H2B hetero-dimers, (CENP-A/H4)_2_ hetero-tetramers and DNA were mixed with a molar ratio of 2:1:1 at high salt concentration (2M NaCl). A gradient dialysis to low salt was performed overnight with a flow rate of 1.5 mL/min using a two-channel peristaltic pump as described in [39]. Assembled nucleosomes were then uniquely positioned on the DNA by a thermal shift for 2h at 55 °C. CENP-A nucleosome and CENP-C^CR^ were complexed by adding 2.2 moles of CENP-C^CR^ to each mole of CENP-A nucleosome. The complex quality was controlled on a 5% native PAGE gel.

### Binding experiments

2.4 μM of CENP-A and H3 nucleosomes assembled on 601 (145 bp) DNA were mixed with different amounts of CENPC^CR^, CENP-C^motif^ or CENP-C^CR-short^ and incubated for 1 hour at 4°C. Complex formation was verified on a 5% native PAGE gel.

### Competition experiments

2.4 μM of the CENP-A/CENP-C^motif^ complex was mixed with different amounts of CENP-C^CR^, and the competition between the two domains was tested using 5% native PAGE gel. 2.4 μM of CENP-A/CENP-C^CR^ complex was mixed with different amounts of CENP-C^motif^ and CENP-C^CR-short^. The competition was then followed using 5% native PAGE gel.

### Micrococcal nuclease digestion

2 μg of nucleosomes were incubated with 2 Kunitz units of Micrococcal nuclease (NEB) in buffer containing 10 mM Tris HCl pH 7.5, 3 mM CaCl_2_ and 1 mM DTT at room temperature. Reactions were quenched at different time points (2, 5, 7, 10 and 20 minutes) by the addition of 250 μL of PB buffer (Qiagen QIAquick PCR Purification Kit) supplemented with 10 mM of EGTA. DNA from each sample was purified with the QIAquick PCR purification kit, and the extent of DNA digestion was quantified by 2100 Bioanalyzer (Agilent). All experiments were done in triplicates.

### CryoEM grid preparation and data collection and processing

CENP-A nucleosome and CENP-A/CENP-C^CR^ complex were prepared as described above. 3 μl of the sample (1-1.2 mg/ml) was applied to freshly glow-discharged Quantifoil R2/1 holey carbon grid. After 3 s blotting time, grids were plunge-frozen in liquid ethane using a FEI Vitrobot automatic plunge freezer. Humidity in the chamber was kept at 95%.

Electron micrographs were recorded on FEI Titan Krios at 300 kV with a Gatan Summit K2 electron detector (∼4700 micrographs) (Cryo-EM facility at MPI for Biochemistry Martinsried, Germany). The image pixel size was 0.65 Å per pixel on the object scale. Data were collected in a defocus range of 7 000 – 30 000 Å with a total exposure of 100 e/Å^2^. 50 frames were collected and aligned with the Unblur software package using a dose filter [40].

Several thousand particles were manually picked and carefully cleaned in Relion to remove inconsistent particles. The resulting useful particles were then used for semi-automatic and automatic particle picking in Relion. The contrast transfer function parameters were determined using CTFFIND4 [41]. The 2D class averages were generated with the Relion software package [42]. Inconsistent class averages were removed from further data analysis. The 3D refinements and classifications were subsequently done in Relion. All final refinements were done using the auto refine option (Relion). The initial reference was filtered to 60 Å and C1 symmetry was applied during refinements for all classes. Particles were split into two datasets and refined independently and the resolution was determined using the 0.143 cut-off (Relion auto refine option). Local resolution was determined with Resmap. All maps were filtered to local resolution using Relion with a B-factor determined by Relion.

### Model building

The model was built in Coot [43] and refined using Phenix real_space_refine [44]. Figures are prepared with Chimera [45].

## Supporting information

Combined supplementary material

## Data availability

Data will be deposited in the EMDB and PDB data banks upon publication.

## Acknowledgements

N.S. and A.A. supported by Centre for Molecular Medicine Norway (NCMM), Department of Chemistry at University of Oslo and Norwegian Research Council, grant # 263195. S.B. and M.H. are supported by St. Jude Children’s Research Hospital, the American Lebanese Syrian Associated Charities and ERC-smallRNAhet-309584. We would like to thank Elena Conti and the cryo-EM facility at Max Planck Institute for Biochemistry in Martinsried for access to cryo-EM microscopes. Without their support this work would not be possible. We also thank Dario Segura-Peña (UiO, Norway) and Lars Jansen (Oxford, UK) for critical reading of the manuscript, and our families for daily support.

## Authors contributions

N.S. and M.H. conceived and supervised the project. A.A. carried out mutagenesis, protein expression, purification and assembled nucleosomes and CENP-C complexes and did MNase experiments and analysis. S.B. prepared grids for cryo-EM. S.B., I.S. and M.H. collected cryoEM data. S.B. and M.H. processed cryo-EM data. A.A., N.S., S.B. and M.H. analyzed data and built the structure and A.A. refined the final models. A.A. and N.S. wrote the manuscript and all authors commented on it.

## EV Figure legends

**Figure EV1. CENP-A nucleosome structural features.**

**A.** Virtual gels from Bioanalyzer showing DNA digestion for CENP-A and H3 nucleosomes on three different DNA templates.

**B.** (Top) Sequence overlay of the N-terminus of H3 and CENP-A, indicating swapped sequences used in CENP-A^H3(N-tail)^ and H3^CENP-A(N-tail)^ constructs. (Bottom) Virtual gels of MNase digestion for CENP-A^H3(N-tail)^ and H3^CENP-A(N-tail)^ nucleosomes assembled on 601 DNA.

**C.** Representative cryoEM map of the CENP-A nucleosome, illustrating quality of map and model fitting.

**D.** EM density of CENP-A specific features on the nucleosome: αN helix (2 different sides), C-terminal tail and RG-loop.

**Figure EV2. CryoEM analysis of CENP-A nucleosome.**

**A.** Representative cryo-EM raw micrograph.

**B.** Subset of selected 2D class averages.

**C.** Euler angle distribution of particles used in the final 3D reconstruction.

**D.** Fourier shell correlation (FSC) curves of the final density map (CENP-A high resolution).

**E.** Local resolution of the final 3D density map.

**F.** Particles used for the high-resolution CENP-A map were further classified for DNA entry/exit site in order to highlight differences at this part of the nucleosome. Gray map (Class 2) has loosest DNA wrap and green map (Class 4) has tightest DNA wrap. The blue map presents particles that where in-between two extreme conformations.

**Figure EV3. CENP-C**^**CR-short**^ **does not bind to the H3 nucleosome.** Native PAGE gel stained with EtBr probing binding of CENP-C^CR-short^ to CENP-A and H3 nucleosome. Lane 1: CENP-A nucleosome. Lanes 3-5: Increasing amounts of CENP-C^CR-short^ are added to CENP-A nucleosome. Generation of sharp band with slower mobility indicates formation of a specific complex. Lane 5: H3 nucleosome. Lanes 6-8: Increasing amounts of CENP-C^CR^ are added to the H3 nucleosome. Band corresponding to H3 nucleosome is not changing mobility, indicating absence of CENP-C^CR-short^/H3 nucleosome complex formation. Interestingly, absence of residues 426-500 in CENP-C^CR-short^ vs. CENP-C^CR^ prevents non-specific binding of H3 nucleosome (Figure 2B). 2.4 μM nucleosomes are used in all conditions.

**Figure EV4. CENP-A nucleosome/CENP-C**^**CR**^ **complex structure.**

**A.** Representative cryoEM densities showing fitted model for DNA and each of the histones (left), arginine core and hydrophobic regions of CENP-C^CR^ (right).

**B.** Space filling model of nucleosome (histone core – grey, DNA - blue), showing a hydrophobic groove (green) on the nucleosome formed by H2A^L108^, CENP-A^L135^, CENP-A^L139^ and H4^V60^. CENP-C^CR^ is shown as a purple coil with hydrophobic sidechains in stick representation.

**Figure EV5. CryoEM analysis of CENP-A/CENP-C**^**CR**^ **complex.**

**A.** Representative cryoEM raw micrograph.

**B.** Subset of selected 2D class averages.

**C.** Euler angle distribution of particles used in the final 3D reconstruction for CENP-A/CENP-C^CR^ high-resolution and CENP-A/CENP-C^CR^ complex enriched for CENP-C^CR^.

**D.** Fourier shell correlation (FSC) curves of the final density map for CENP-A/CENP-C^CR^ high-resolution and CENP-A/CENP-C^CR^ enriched for CENP-C^CR^.

**E.** Local resolution of the final 3D density maps.

**F.** Thefirst particles were sorted for high-resolution, and this map was used for initial model building. A map enriched in CENP-C^CR^ was generated to increase map quality around CENP-C^CR^. Particles used for the later map were further classified for DNA entry/exit site in order to highlight extend of DNA unwrapping. The gray map (Class 2) has the loosest DNA wrap and the green map (Class 4) has tightest DNA wrap. The blue map presents particles that where in-between two extreme conformations.

**Figure EV6. Conformational changes on the CENP-A nucleosome upon CENP-C**^**CR**^ **binding. A-C.** Virtual gels for MNase digestion.

**A.** MNase digestion of the CENP-A nucleosome alone and in complex with CENP-C^CR^ or CENP-C^motif^ (also in Figure 4A)

**B.** MNase digestion of the CENP-A nucleosome assembled with H2A lacking 110-130 residues alone (top) or in complex with CENP-C^CR^ (bottom), showing similar magnitude of digestion.

**C.** (Top) MNase digestion of the H3^ΔC-tail H2A^, indicating that removal of H2A^110-130^ does not have an effect on the DNA digestion speed in the context of H3 nucleosome. (Middle) MNase digestion of CENP-A^H3(N-tail),^ ΔC-tail H2A and H3^CENP-A(N-tail),^ ΔC-tail H2A, indicating that removal of H2A^110-130^ does not have an effect on the DNA digestion speed in the context of CENP-A^H3(N-tail)^ nucleosome, but digestion is slightly increased in the context of H3^CENP-A(N-tail)^. (Bottom) MNase digestion of the CENP-A^H3(N-tail)^ is unaffected with CENP-C^CR^ binding independently of the presence of H2A^110-130^.

**D.** Interactions between H2A C-terminal tail and CENP-C^CR^.

## References

1. McKinley KL, Cheeseman IM (2016) The molecular basis for centromere identity and function. Nat Rev Mol Cell Biol 17: 16–29.

2. Potapova T, Gorbsky GJ (2017) The Consequences of Chromosome Segregation Errors in Mitosis and Meiosis. Biology 6: 12.

3. Sekulic N, Black BE (2012) Molecular underpinnings of centromere identity and maintenance. Trends Biochem Sci 37: 220–229.

4. Carroll CW, Milks KJ, Straight AF (2010) Dual recognition of CENP-A nucleosomes is required for centromere assembly. J Cell Biol 189: 1143–1155.

5. Sekulic N, Bassett EA, Rogers DJ, Black BE (2010) The structure of (CENP-A-H4)2 reveals physical features that mark centromeres. Nature 467: 347–351.

6. Tachiwana H, Kagawa W, Shiga T, Osakabe A, Miya Y, Saito K, Hayashi-Takanaka Y, Oda T, Sato M, Park S-Y, et al. (2011) Crystal structure of the human centromeric nucleosome containing CENP-A. Nature 476: 232–235.

7. Pentakota S, Zhou K, Smith C, Maffini S, Petrovic A, Morgan GP, Weir JR, Vetter IR, Musacchio A, Luger K (2017) Decoding the centromeric nucleosome through CENP-N. eLife 6:.

8. Chittori S, Hong J, Saunders H, Feng H, Ghirlando R, Kelly AE, Bai Y, Subramaniam S (2018) Structural mechanisms of centromeric nucleosome recognition by the kinetochore protein CENP-N. Science 359: 339–343.

9. Tian T, Li X, Liu Y, Wang C, Liu X, Bi G, Zhang X, Yao X, Zhou ZH, Zang J (2018) Molecular basis for CENP-N recognition of CENP-A nucleosome on the human kinetochore. Cell Res 28: 374–378.

10. Musacchio A, Desai A (2017) A Molecular View of Kinetochore Assembly and Function. Biology 6: 5.

11. Milks KJ, Moree B, Straight AF (2009) Dissection of CENP-C–directed Centromere and Kinetochore Assembly. Mol Biol Cell 20: 4246–4255.

12. Song K, Gronemeyer B, Lu W, Eugster E, Tomkiel JE (2002) Mutational analysis of the central centromere targeting domain of human centromere protein C, (CENP-C). Exp Cell Res 275: 81–91.

13. Guo LY, Allu PK, Zandarashvili L, McKinley KL, Sekulic N, Dawicki-McKenna JM, Fachinetti D, Logsdon GA, Jamiolkowski RM, Cleveland DW, et al. (2017) Centromeres are maintained by fastening CENP-A to DNA and directing an arginine anchor-dependent nucleosome transition. Nat Commun 8: 15775.

14. Kato H, Jiang J, Zhou B-R, Rozendaal M, Feng H, Ghirlando R, Xiao TS, Straight AF, Bai Y (2013) A Conserved Mechanism for Centromeric Nucleosome Recognition by Centromere Protein CENP-C. Science 340: 1110–1113.

15. Black BE, Cleveland DW (2011) Epigenetic Centromere Propagation and the Nature of CENP-A Nucleosomes. Cell 144: 471–479.

16. Hasson D, Panchenko T, Salimian KJ, Salman MU, Sekulic N, Alonso A, Warburton PE, Black BE (2013) The octamer is the major form of CENP-A nucleosomes at human centromeres. Nat Struct Mol Biol 20: 687–695.

17. Nechemia-Arbely Y, Fachinetti D, Miga KH, Sekulic N, Soni GV, Kim DH, Wong AK, Lee AY, Nguyen K, Dekker C, et al. (2017) Human centromeric CENP-A chromatin is a homotypic, octameric nucleosome at all cell cycle points. J Cell Biol 216: 607–621.

18. Roulland Y, Ouararhni K, Naidenov M, Ramos L, Shuaib M, Syed SH, Lone IN, Boopathi R, Fontaine E, Papai G, et al. (2016) The Flexible Ends of CENP-A Nucleosome Are Required for Mitotic Fidelity. Mol Cell 63: 674–685.

19. Condee Silva N, Black BE, Sivolob A, Filipski J, Cleveland DW, Prunell A (2007) CENP-A-containing Nucleosomes: Easier Disassembly versus Exclusive Centromeric Localization. J Mol Biol 370: 555–573.

20. Zhou B-R, Yadav KNS, Borgnia M, Hong J, Cao B, Olins AL, Olins DE, Bai Y, Zhang P (2019) Atomic resolution cryo-EM structure of a native-like CENP-A nucleosome aided by an antibody fragment. Nat Commun 10: 2301.

21. Lowary PT, Widom J (1998) New DNA sequence rules for high affinity binding to histone octamer and sequence-directed nucleosome positioning1. J Mol Biol 276: 19–42.

22. Masumoto H, Masukata H, Muro Y, Nozaki N, Okazaki T (1989) A human centromere antigen (CENP-B) interacts with a short specific sequence in alphoid DNA, a human centromeric satellite. J Cell Biol 109: 1963–1973.

23. Panchenko T, Sorensen TC, Woodcock CL, Kan Z, Wood S, Resch MG, Luger K, Englander SW, Hansen JC, Black BE (2011) Replacement of histone H3 with CENP-A directs global nucleosome array condensation and loosening of nucleosome superhelical termini. Proc Natl Acad Sci 108: 16588–16593.

24. Bilokapic S, Strauss M, Halic M (2018) Histone octamer rearranges to adapt to DNA unwrapping. Nat Struct Mol Biol 25: 101.

25. Carroll CW, Silva MCC, Godek KM, Jansen LET, Straight AF (2009) Centromere assembly requires the direct recognition of CENP-A nucleosomes by CENP-N. Nat Cell Biol 11: 896–902.

26. McGinty RK, Tan S (2016) Recognition of the nucleosome by chromatin factors and enzymes. Curr Opin Struct Biol 37: 54–61.

27. Falk SJ, Guo LY, Sekulic N, Smoak EM, Mani T, Logsdon GA, Gupta K, Jansen LET, Duyne GDV, Vinogradov SA, et al. (2015) CENP-C reshapes and stabilizes CENP-A nucleosomes at the centromere. Science 348: 699–703.

28. Eickbush TH, Godfrey JE, Elia MC, Moudrianakis EN (1988) H2a-specific proteolysis as a unique probe in the analysis of the histone octamer. J Biol Chem 263: 18972–18978.

29. Vogler C, Huber C, Waldmann T, Ettig R, Braun L, Izzo A, Daujat S, Chassignet I, Lopez-Contreras AJ, Fernandez-Capetillo O, et al. (2010) Histone H2A C-Terminus Regulates Chromatin Dynamics, Remodeling, and Histone H1 Binding. PLOS Genet 6: e1001234.

30. Biswas M, Voltz K, Smith JC, Langowski J (2011) Role of Histone Tails in Structural Stability of the Nucleosome. PLOS Comput Biol 7: e1002279.

31. Hori T, Shang W-H, Toyoda A, Misu S, Monma N, Ikeo K, Molina O, Vargiu G, Fujiyama A, Kimura H, et al. (2014) Histone H4 Lys 20 Monomethylation of the CENP-A Nucleosome Is Essential for Kinetochore Assembly. Dev Cell 29: 740–749.

32. Arimura Y, Tachiwana H, Takagi H, Hori T, Kimura H, Fukagawa T, Kurumizaka H (2019) The CENP-A centromere targeting domain facilitates H4K20 monomethylation in the nucleosome by structural polymorphism. Nat Commun 10: 576.

33. Allu PK, Dawicki-McKenna JM, Eeuwen TV, Slavin M, Braitbard M, Xu C, Kalisman N, Murakami K, Black BE (2019) Structure of the Human Core Centromeric Nucleosome Complex. Curr Biol 0:

34. Bönisch C, Hake SB (2012) Histone H2A variants in nucleosomes and chromatin: more or less stable? Nucleic Acids Res 40: 10719–10741.

35. Bodor DL, Mata JF, Sergeev M, David AF, Salimian KJ, Panchenko T, Cleveland DW, Black BE, Shah JV, Jansen LE (2014) The quantitative architecture of centromeric chromatin. eLife 3: e02137.

36. Ross JE, Woodlief KS, Sullivan BA (2016) Inheritance of the CENP-A chromatin domain is spatially and temporally constrained at human centromeres. Epigenetics Chromatin 9: 20.

37. Sekulic N, Black BE (2016) Preparation of Recombinant Centromeric Nucleosomes and Formation of Complexes with Nonhistone Centromere Proteins. Methods Enzymol 573: 67–96.

38. Dyer PN, Edayathumangalam RS, White CL, Bao Y, Chakravarthy S, Muthurajan UM, Luger K (2003) Reconstitution of Nucleosome Core Particles from Recombinant Histones and DNA. In, Methods in Enzymology pp 23–44. Academic Press.

39. Luger K, Rechsteiner TJ, Richmond TJ (1999) Preparation of nucleosome core particle from recombinant histones. In, Methods in Enzymology pp 3–19. Academic Press.

40. Grant T, Grigorieff N (2015) Measuring the optimal exposure for single particle cryo-EM using a 2.6 Å reconstruction of rotavirus VP6. eLife 4: e06980.

41. Rohou A, Grigorieff N (2015) CTFFIND4: Fast and accurate defocus estimation from electron micrographs. J Struct Biol 192: 216–221.

42. Scheres SHW (2012) RELION: Implementation of a Bayesian approach to cryo-EM structure determination. J Struct Biol 180: 519–530.

43. Emsley P, Lohkamp B, Scott WG, Cowtan K (2010) Features and development of Coot. Acta Crystallogr D Biol Crystallogr 66: 486–501.

44. Adams PD, Afonine PV, Bunkóczi G, Chen VB, Davis IW, Echols N, Headd JJ, Hung L-W, Kapral GJ, Grosse-Kunstleve RW, et al. (2010) PHENIX: a comprehensive Python-based system for macromolecular structure solution. Acta Crystallogr D Biol Crystallogr 66: 213–221.

45. Pettersen EF, Goddard TD, Huang CC, Couch GS, Greenblatt DM, Meng EC, Ferrin TE (2004) UCSF Chimera--a visualization system for exploratory research and analysis. J Comput Chem 25: 1605–1612.

