## Supplementary material for "CENP-C unwraps the CENP-A nucleosome through the H2A C-terminal tail": Combined supplementary material

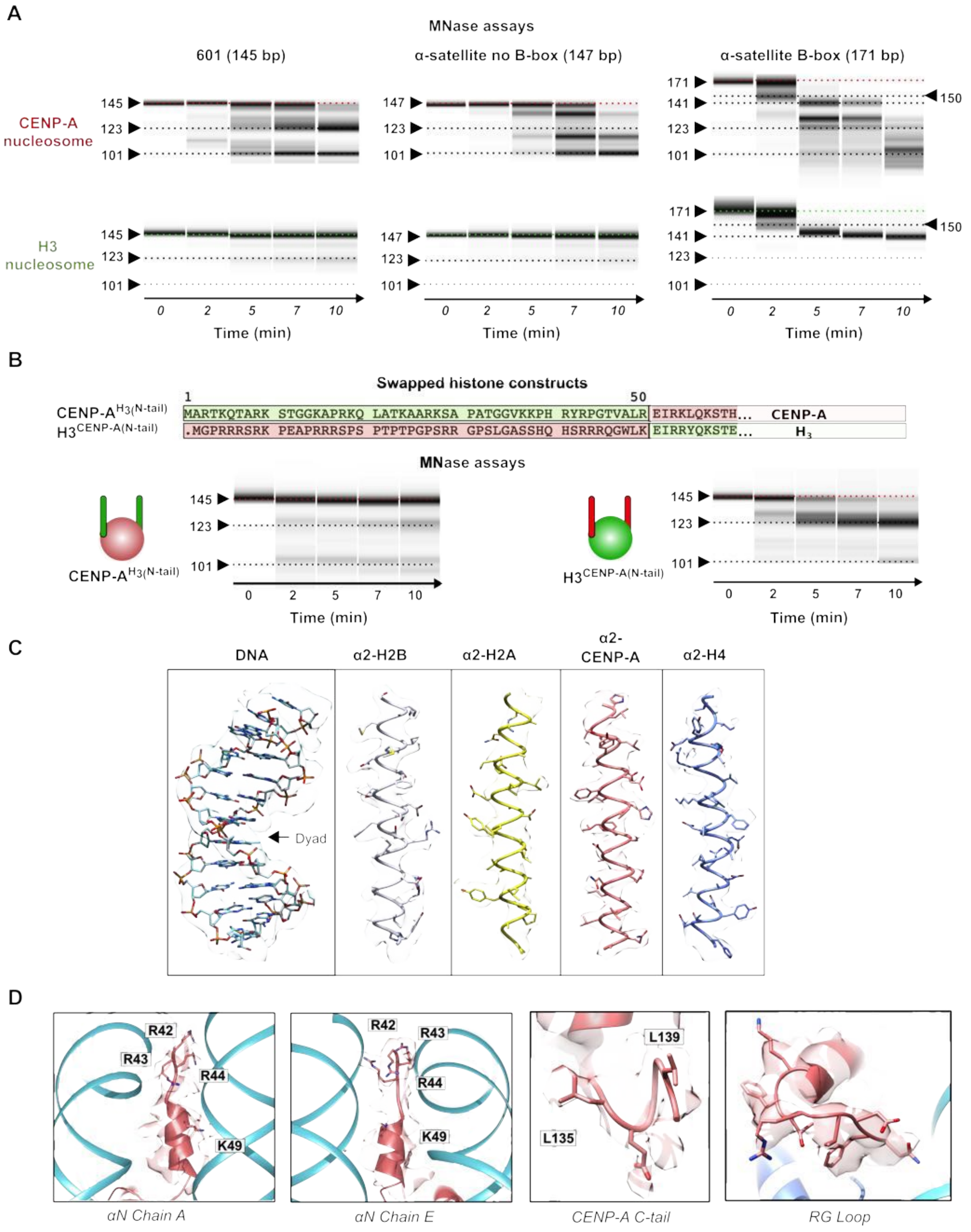

A
MNase assays
601 (145 bp)
α-satellite no B-box (147 bp)
α-satellite B-box (171 bp)
CENP-A nucleosome
H3 nucleosome
145
123
101
147
171
141
150
0 2 5 7 10
Time (min)
B
Swapped histone constructs
1
50
CENP-A^H3(N-tail)
H3^CENP-A(N-tail)
MARTKQTARK STGGKAPRKQ LATKAARKSA PATGGVKKPH RYRPGTVALR EIRKLQKSTH... CENP-A
.MGPRRRSRK PEAPRRRSPS PTPTPGPSRR GPSLGASSHQ HSRRRQGWLK EIRRYQKSTE... H3
MNase assays
C
DNA
α2-H2B
α2-H2A
α2-CENP-A
α2-H4
Dyad
D
R42
R43
R44
K49
L139
L135
αN Chain A
αN Chain E
CENP-A C-tail
RG Loop

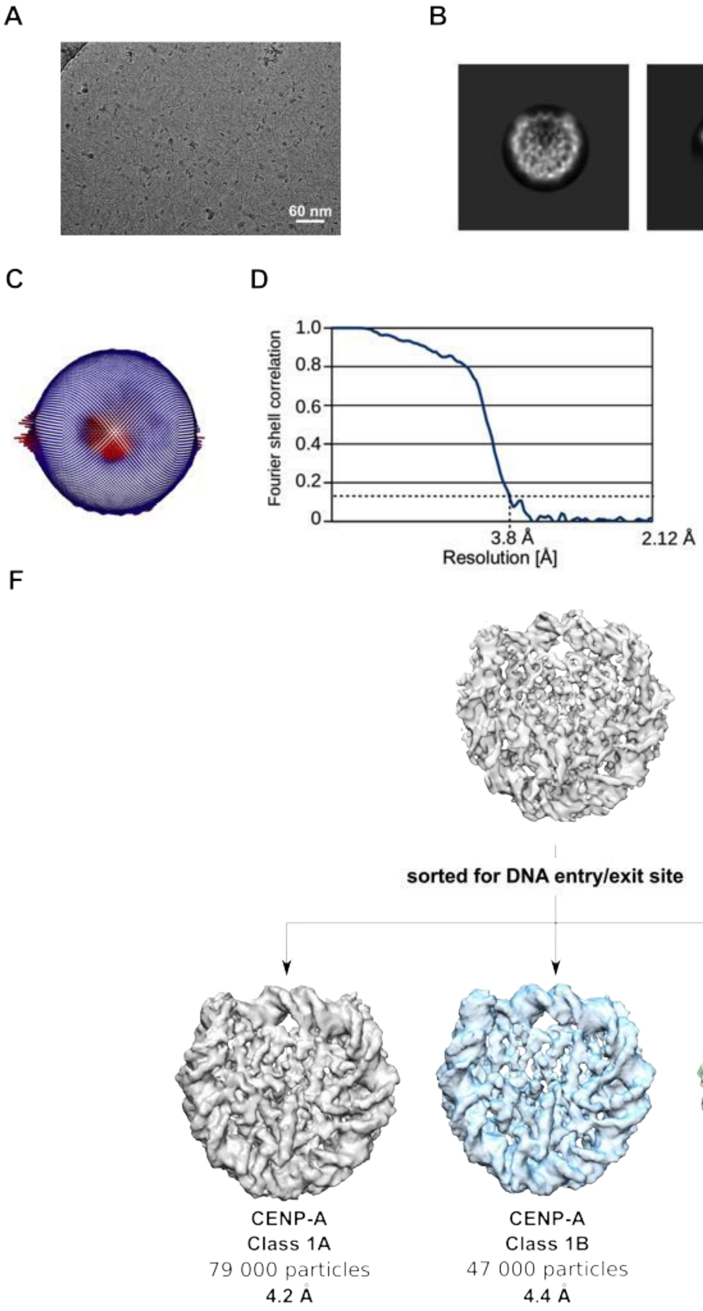
A
60 nm
B
C
D
1.0
0.8
0.6
0.4
0.2
0
Fourier shell correlation
3.8 Å
2.12 Å
Resolution [Å]
E
2.8 Å
3.2 Å
3.6 Å
4.0 Å
4.4 Å
F
CENP-A
high-resolution
Class 1
230 000 particles
3.8 Å
sorted for DNA entry/exit site
CENP-A
Class 1A
79 000 particles
4.2 Å
CENP-A
Class 1B
47 000 particles
4.4 Å
CENP-A
Class 1C
104 000 particles
4.2 Å

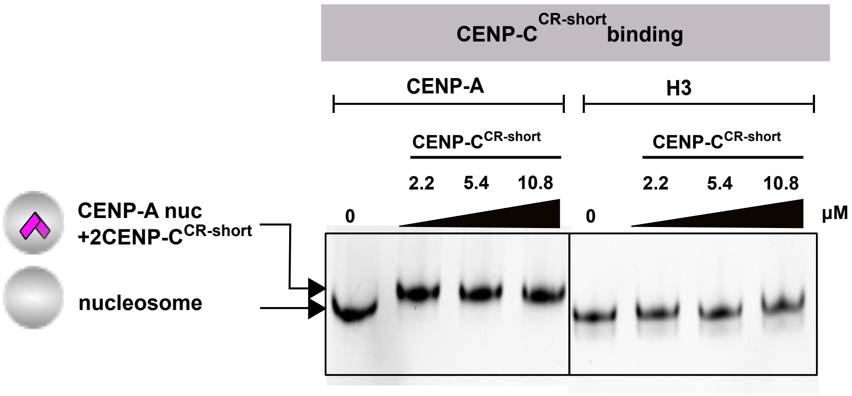
CENP-C CR-short binding
CENP-A
H3
CENP-C CR-short
CENP-C CR-short
2.2 5.4 10.8
2.2 5.4 10.8
0
0
μM
CENP-A nuc +2CENP-C CR-short
nucleosome

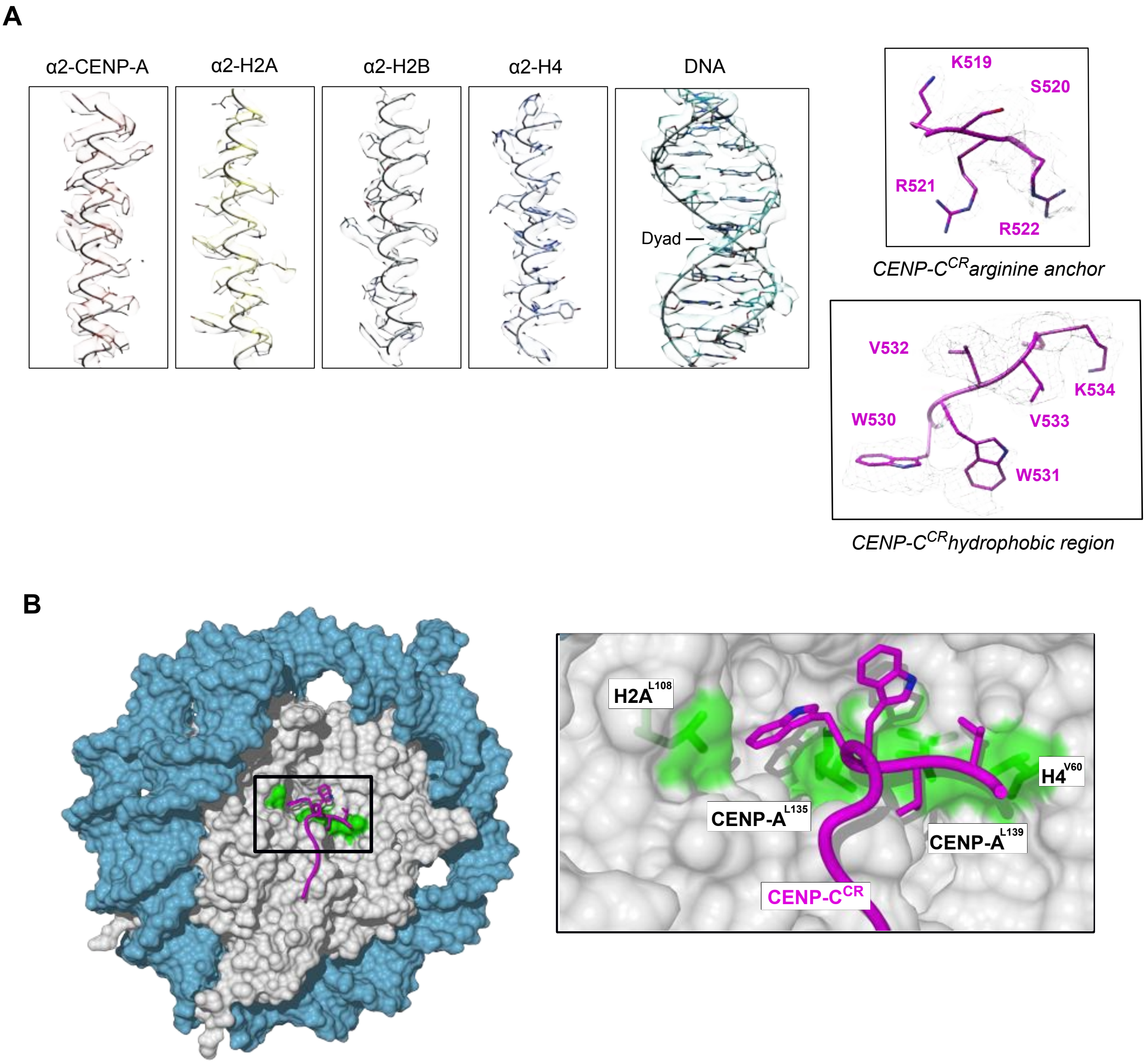
A
α2-CENP-A
α2-H2A
α2-H2B
α2-H4
DNA
Dyad
K519
S520
R521
R522
CENP-C^CR arginine anchor
V532
K534
W530
V533
W531
CENP-C^CR hydrophobic region
B
H2A^L108
H4^V60
CENP-A^L135
CENP-A^L139
CENP-C^CR

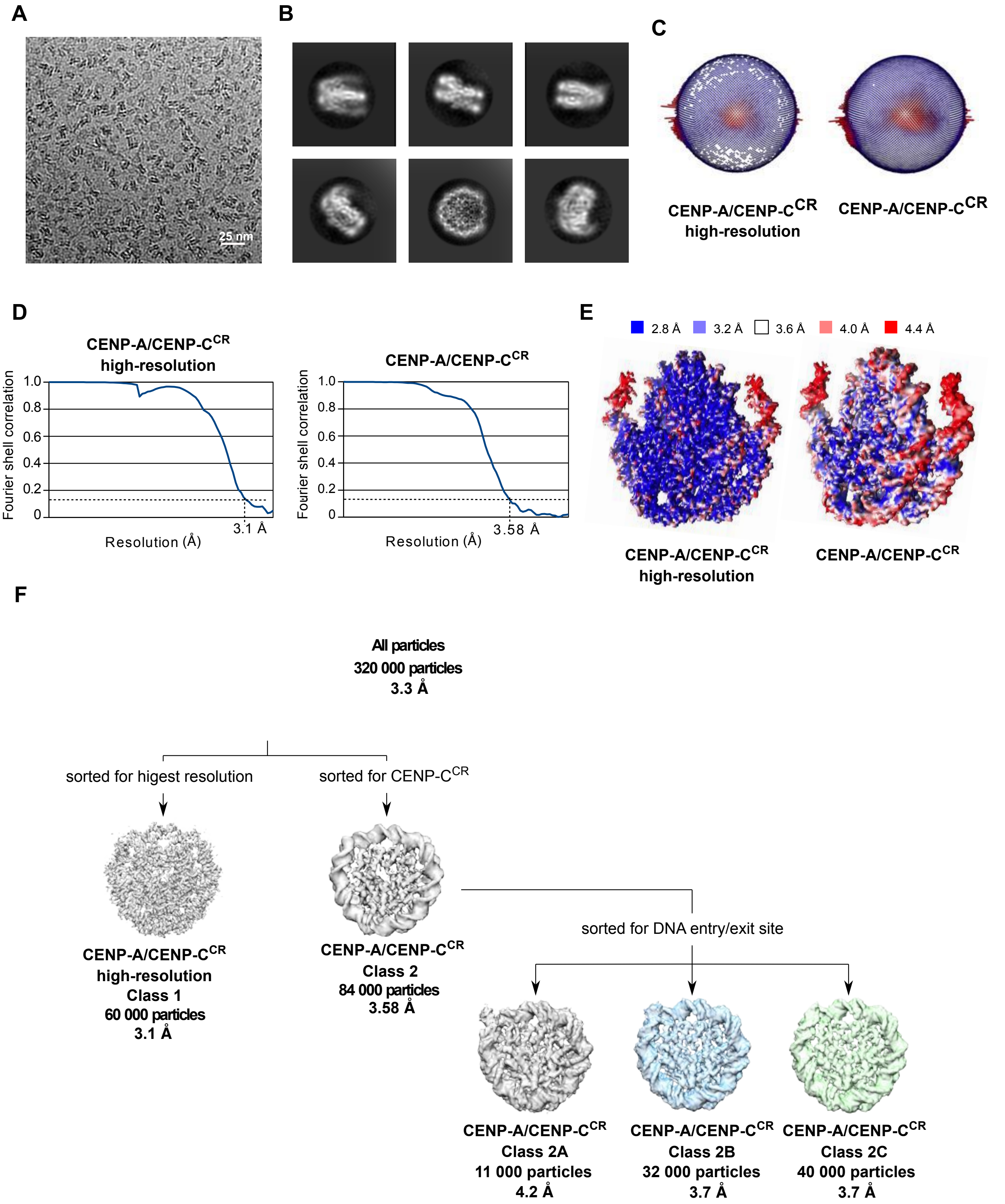
A
25 nm
B
C
CENP-A/CENP-C$^{CR}$ high-resolution
CENP-A/CENP-C$^{CR}$
D
CENP-A/CENP-C$^{CR}$ high-resolution
Fourier shell correlation
1.0
0.8
0.6
0.4
0.2
0
3.1 Å
Resolution (Å)
CENP-A/CENP-C$^{CR}$
3.58 Å
E
2.8 Å
3.2 Å
3.6 Å
4.0 Å
4.4 Å
F
All particles
320 000 particles
3.3 Å
sorted for higest resolution
sorted for CENP-C$^{CR}$
CENP-A/CENP-C$^{CR}$ high-resolution Class 1 60 000 particles 3.1 Å
CENP-A/CENP-C$^{CR}$ Class 2 84 000 particles 3.58 Å
sorted for DNA entry/exit site
CENP-A/CENP-C$^{CR}$ Class 2A 11 000 particles 4.2 Å
CENP-A/CENP-C$^{CR}$ Class 2B 32 000 particles 3.7 Å
CENP-A/CENP-C$^{CR}$ Class 2C 40 000 particles 3.7 Å

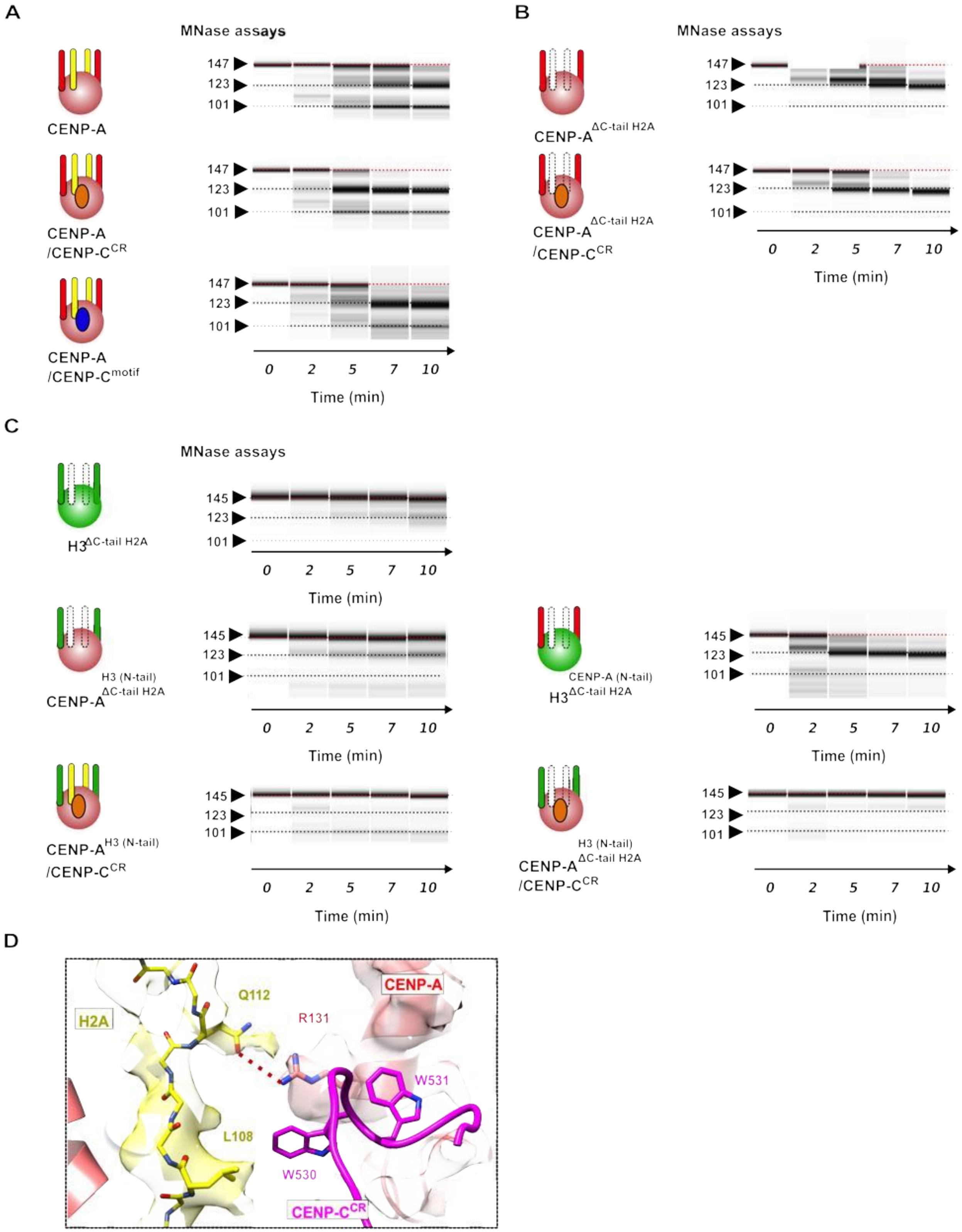
A
MNase assays
CENP-A
CENP-A /CENP-C^CR
CENP-A /CENP-C^motif
147
123
101
0 2 5 7 10
Time (min)
B
MNase assays
CENP-A^ΔC-tail H2A
CENP-A^ΔC-tail H2A /CENP-C^CR
C
MNase assays
H3^ΔC-tail H2A
145
CENP-A^H3 (N-tail) ΔC-tail H2A
H3^CENP-A (N-tail) ΔC-tail H2A
CENP-A^H3 (N-tail) /CENP-C^CR
CENP-A^H3 (N-tail) ΔC-tail H2A /CENP-C^CR
D
H2A
Q112
R131
CENP-A
W531
L108
W530
CENP-C^CR

**Appendix Table 1: Cryo-EM data collection, refinement and validation statistics for CENP-A nucleosome**

| | Class 1<br>EMD- 10151<br>6SE0 | Class 1A<br>EMD- 10157 | Class 1B<br>EMD- 10158 | Class 1C<br>EMD- 10159 |
|---|---|---|---|---|
| **Data collection and processing** | | | | |
| Magnification | | | | |
| Voltage (kV) | 200 | 200 | 200 | 200 |
| Electron exposure (e–/Å$^2$) | 80 | 80 | 80 | 80 |
| Defocus range (μm) | -0.7 – -3.0 | -0.7– -3.0 | - 0.7 – -3.0 | - 0.7 – -3.0 |
| Pixel size (Å) | 1.06 | 1.06 | 1.06 | 1.06 |
| Symmetry imposed | C1 | C1 | C1 | C1 |
| Initial particle images (no.) | ~ 230 000 | ~ 230 000 | ~ 230 000 | ~ 230 000 |
| Final particle images (no.) | 170 000 | 79 000 | 47 000 | 104 000 |
| Map resolution (Å)<br>FSC threshold | 3.8 | 4.2 | 4.4 | 4.2 |
| Map resolution range (Å) | 4-6.0 | 4-6.0 | 4-6.0 | 4-6.0 |
| **Refinement** | | | | |
| Initial model used | 6E0C | | | |
| Model resolution (Å)<br>FSC threshold | 3.8 | | | |
| Model resolution range (Å) | 235-3.8 | | | |
| Map sharpening *B* factor (Å$^2$) | -100 | | | |
| Model composition<br>Nonhydrogen atoms<br>Protein residues<br>Ligands | <br>12002<br>765<br>0 | | | |
| R.m.s. deviations<br>Bond lengths (Å)<br>Bond angles (°) | <br>0.008<br>1.01 | | | |
| Validation<br>MolProbity score.<br>Clashscore<br>Poor rotamers (%) | <br>1.17<br>3.83<br>0 | | | |
| Ramachandran plot<br>Favored (%)<br>Allowed (%)<br>Disallowed (%) | <br>98.8<br>1.17<br>0 | | | |

**Appendix Table 2: Cryo-EM data collection, refinement and validation statistics for CENP-A nucleosome/ CENP-C central region complex**

| | Class 1<br>EMD- 10155<br>6SEG | Class 2<br>EMD-10152<br>6SE6 | Class 2A<br>EMD- 10153<br>6SEE | Class 2B<br>EMD- 10156 | Class 2C<br>EMD- 10154<br>6SEF |
|---|---|---|---|---|---|
| **Data collection and processing** | | | | | |
| Magnification | | | | | |
| Voltage (kV) | 300 | 300 | 300 | 300 | 300 |
| Electron exposure (e–/Å$^2$) | 100 | 100 | 100 | 100 | 100 |
| Defocus range (μm) | -0.7 – -3.0 | -0.7– -3.0 | - 0.7 – -3.0 | -0.7 – -3.0 | -0.7 – -3.0 |
| Pixel size (Å) | 0.65 (1.3) | 0.65 (1.3) | 0.65 (1.3) | 0.65 (1.3) | 0.65 (1.3) |
| Symmetry imposed | C1 | C1 | C1 | C1 | C1 |
| Initial particle images (no.) | ~ 320 000 | ~ 320 000 | ~ 84 000 | ~ 84 000 | ~ 84 000 |
| Final particle images (no.) | 60 000 | 84 000 | 11 000 | 32 000 | 40 000 |
| Map resolution (Å)<br>FSC threshold | 3.1 | 3.5 | 4.2 | 3.7 | 3.7 |
| Map resolution range (Å) | 2.6-5.0 | 3.3 – 5.0 | 4.0 – 5.0 | 3.5-5.0 | 3.5-5.0 |
| **Refinement** | | | | | |
| Initial model used | 6BUZ | Class 1 | Class 1 | | Class 1 |
| Model resolution (Å)<br>FSC threshold | 3.1 | 3.6 | 4.2 | | 3.7 |
| Model resolution range (Å) | 235-2.9 | 235-2.9 | 235-3.6 | | 235-3.8 |
| Map sharpening *B* factor (Å$^2$) | -100 | -100 | -100 | | -100 |
| Model composition | | | | | |
| Nonhydrogen atoms | 11690 | 12068 | 11789 | | 11759 |
| Protein residues | 728 | 771 | 738 | | 733 |
| Ligands | 0 | 0 | 0 | | 0 |
| R.m.s. deviations | | | | | |
| Bond lengths (Å) | 0.006 | 0.010 | 0.008 | | 0.008 |
| Bond angles (°) | 0.849 | 0.984 | 0.940 | | 0.855 |
| Validation | | | | | |
| MolProbity score. | 1.04 | 1.14 | 1.39 | | 1.42 |
| Clashscore | 2.54 | 3.44 | 4.25 | | 5.11 |
| Poor rotamers (%) | 0 | 0 | 0 | | 0 |
| Ramachandran plot | | | | | |
| Favored (%) | 98.74 | 98..67 | 96.95 | | 97.21 |
| Allowed (%) | 1.26 | 1.33 | 3.05 | | 2.79 |
| Disallowed (%) | 0 | 0 | 0 | | 0 |
